# Micromapping (µMap) of HER2 Across Human Breast Cancers: Photocatalytic Proximity Labeling Identifies Primary Resistance Mechanisms and Functional Interactors

**DOI:** 10.1101/2025.06.10.658685

**Authors:** Steve D. Knutson, Jacob A. Boyer, Danielle C. Morgan, Johannes Großkopf, Joshua D. Rabinowitz, David W. C. MacMillan

## Abstract

Interactions among receptor tyrosine kinases (RTKs), particularly HER2, are critical in driving the growth of certain breast cancer subtypes. However, the mechanisms of HER2 dependency and resistance to targeted therapies remain unclear. Using photocatalytic micromapping (µMap) proximity labeling, we profiled the HER2 interactome across human breast cancer lines. We identify galectin family proteins as uniquely enriched in trastuzumab-resistant models, and show that genetic and pharmacological inhibition of galectins restores trastuzumab sensitivity. Mechanistically, galectin inhibition destabilized HER2 and other RTKs and enhanced antibody-mediated receptor degradation. Galectin inhibition in combination with trastuzumab triggered mitochondrial and endoplasmic reticulum stress pathways, revealing new mechanisms underlying HER2 signaling, dependency, and resistance in breast cancer. We also identified protein tyrosine phosphatase F (PTPRF) as a pan-cancer HER2 interactor, which is broadly upregulated and whose knockdown suppresses proliferation in HER2-low cancers. This work provides an extensive new interactomic resource and underscores the utility of proximity labeling for mapping complex cancer networks and identifying new therapeutic targets.

## INTRODUCTION

Breast cancer remains a leading cause of cancer-related death among women worldwide. Cancer initiation and progression are driven by dysregulated growth signaling networks, including receptor tyrosine kinases (RTKs) and their effectors. HER2, a member of the ErbB receptor tyrosine kinase family, is amplified in approximately 15-20% of breast cancers, defining the HER2+ subtype.^1^ Several targeted therapies have been developed against HER2, including kinase inhibitors lapatinib and tucatinib, monoclonal antibodies trastuzumab (Tz) and pertuzumab, and several antibody-drug conjugates. These treatments have significantly improved patient outcomes, although intrinsic and acquired resistance remains a major challenge.^2,3^ Additionally, while not amplified, HER2 often remains expressed in the HER2-negative (–) setting, where its function and biochemistry are poorly understood.

HER2 signaling is initiated at the plasma membrane and requires dimerization with other ERBB family members such as EGFR, HER3, and HER4 to activate downstream effectors that govern proliferation and survival in cancer.^4^ Other plasma membrane proteins have been proposed as HER2 interactors, but their functions remain unclear. In addition, a comprehensive portrait of the HER2 interactome in breast cancer is unknown.

High-resolution profiling of receptor interactomes can address current limitations in mechanistic understanding and facilitate novel therapeutic intervention.^5, 6^ Traditional biochemical approaches such as immunoprecipitation (IP) have been instrumental in mapping key nodes and individual components in several pathways.^7^ However, IP is problematic for profiling cell-surface interactomes as membrane proteins are often difficult to solubilize without disrupting structure or native interactions. Additionally, interactor abundance can span large dynamic ranges, and IP often struggles to capture transient or spatially restricted interactions, particularly within the dynamically crowded cell-surface environment.

Proximity labeling has emerged as a powerful alternative to traditional IP-based approaches, whereby a catalyst is targeted to a protein-of-interest to label nearby biomolecules.^8, 9^ Our group has recently developed a photocatalytic labeling strategy for “microenvironment or micro-mapping” (µMap),^10^ which employs diazirine-based labeling through activation by a photocatalyst. We initially used this method to map interactions of programmed death-ligand 1 (PD-L1) on T-cells and to study immune synapse associations.^10^ The µMap platform has since been applied in a variety of biological contexts, including chromatin organization,^11^ subcellular RNA localization,^12^ leukocyte and viral receptor interfaces,^13, 14^ and small molecule drug targets and binding sites.^15-17^ Given its utility across these diverse contexts, µMap is ideally suited for resolving HER2-centric interactions, enabling high-resolution phenotyping across tumor subtypes to uncover resistance mechanisms and identify new therapeutic targets.

Here, we employ this technology to map the HER2 interactome in 11 cell lines across breast cancer subtypes, including luminal A (HR+/HER2-) and B (HR+/HER2+), HER2-enriched (HR-/HER2+), Triple-Negative Breast Cancer (TNBC, HR-/HER2-), and normal mammary cells. Our findings reveal protein interactors that are common to HER2+ and HER2– cancers as well as unique signatures underlying dependency and Tz sensitivity. We also identify protein tyrosine phosphatase F (PTPRF) as a cancer-specific interactor and show its functional influence on cell proliferation. These data not only provide a new proteomic resource that describes the HER2 interactome across diverse breast cancer models, but also provides new mechanistic insight regarding HER2 dependency and Tz sensitivity. Together, this work demonstrates the utility of µMap profiling for providing deeper understanding of extracellular receptor signaling networks and, in turn, facilitating the discovery and development of novel targeted cancer therapies.

## RESULTS

### HER2 Interactome Profiling using µMap

To understand the relationship between the HER2 interactome, dependency, and therapeutic response, we identified candidate breast cancer cell lines using CRISPR knock-out loss-of-function data from the Cancer Dependency Map project (DepMap) (**Figure 1A**).^18^ Plotting HER2 expression levels versus dependency across ∼50 breast cancer cell lines illustrates the current clinical challenge that the majority of breast tumors remain untreatable using HER2-targeted therapies. Most cell lines either exhibit low target receptor expression or are HER2-independent and thus have natural resistance to Tz or similar targeted therapies. (**Figure 1A**).

**Figure 1.**
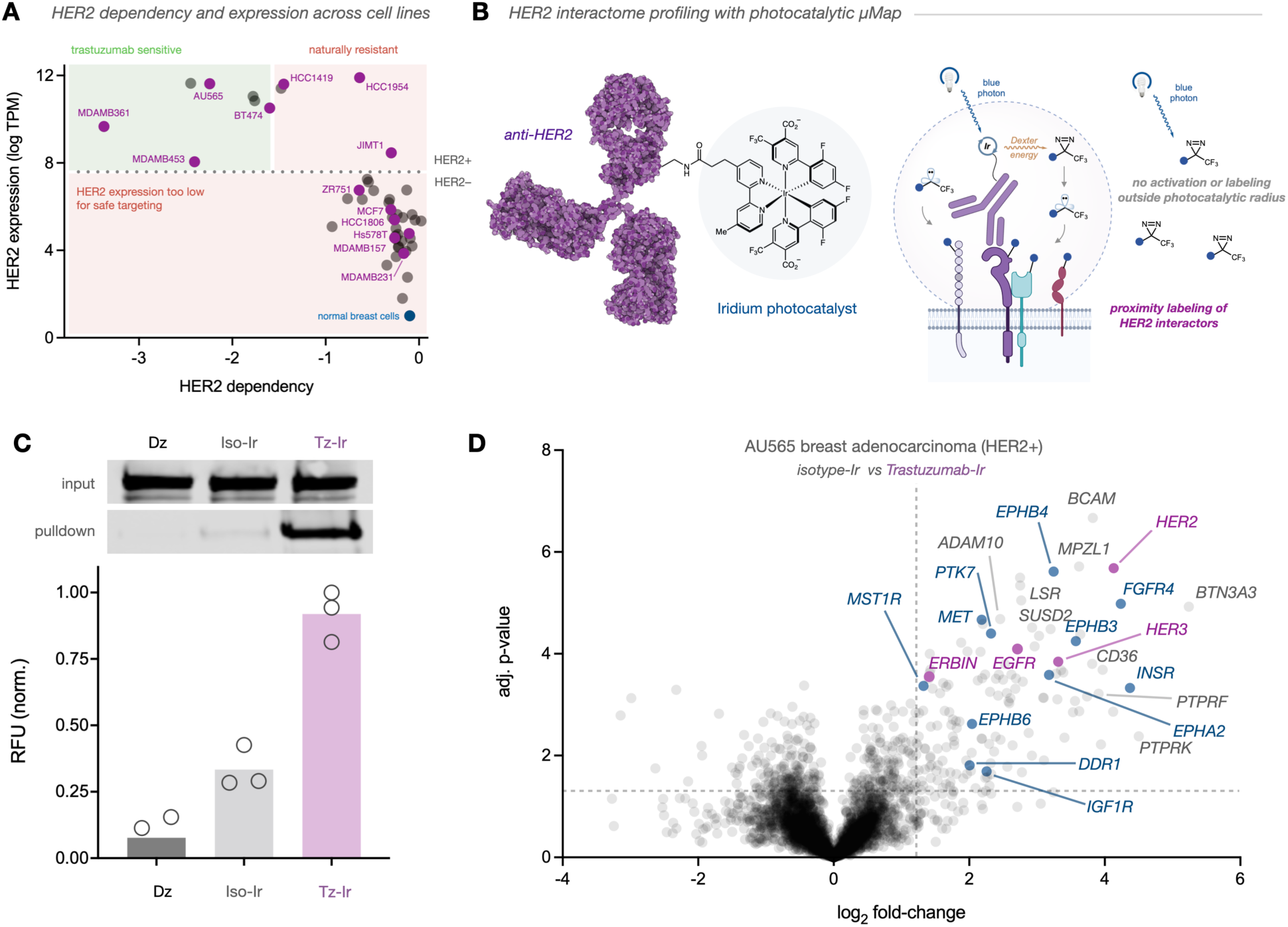
Photocatalytic Mapping of the HER2 Microenvironment. (A) DepMap scatter plot of HER2 expression levels (log TPM) and HER2 dependency scores. (B) Schematic representation of μMap proximity labeling to identify HER2 interactors. (C) Western blot and fluorescent staining evaluation of HER2 photolabeling in AU565 cells targeted with diazirine-biotin alone (Dz) or a photocatalyst-conjugated isotype control (Iso-Ir) or trastuzumab (Tz-Ir). (D) Volcano plot of HER2 µMap proteomic analysis in AU565 breast adenocarcinoma. Several significantly enriched proteins (*p* < 0.05) are highlighted and labeled, including known HER2 interactors (purple) and RTKs (blue).

Spatial interactome profiling with µMap can be accomplished using photocatalyst-conjugated antibodies to label proteins in the immediate vicinity of cell surface targets. Upon visible-light excitation of the photocatalyst, subsequent catalyst activation of diazirine-biotin (Dz) probes to form carbenes ensures that proximal proteins are covalently targeted within a highly precise and narrow radius, thereby enabling streptavidin-based enrichment and subsequent label-free liquid chromatography mass spectrometry (LC−MS) (**Figure 1B**). We first constructed Tz or human IgG isotype control Iridium (Ir) photocatalyst-antibody conjugates and performed targeted photolabeling on the HER2+ AU565 cell line. At the outset, we confirmed the ability of the Tz-Ir conjugate to specifically enrich for HER2—a prerequisite for identifying nearby interactors. Fluorescent staining of live cells treated with either Dz alone, isotype-Ir (Iso-Ir), or Tz-Ir using streptavidin AlexaFluor 555 revealed significant cell-surface labeling in the Tz-Ir sample. Western blot analysis of enriched lysates confirmed robust HER2 labeling and isolation (**Figure 1C**). Following validation, we performed a full µMap workflow on AU565 cells with quantitative LC-MS proteomics of enriched lysates. **Figure 1D** displays a volcano plot of the HER2 proteomic interactome (**Table S1**). In addition to HER2 itself, we also observed an enrichment of other known HER2 interactors, including EGFR (ERBB1), HER3 (ERBB3), and ERBB2 interacting protein (ERBIN) (purple). We also observed significant interaction with other RTKs, including Proto-Oncogene, Receptor Tyrosine Kinase (MET),^19^ EPH Receptors A2 (EPHA2), B3 (EPHB3), B4 (EPHB4), and B6 (EPHB6),^20^ Insulin Receptor (INSR) and Insulin Like Growth Factor 1 Receptor (IGF1R),^21^ and Fibroblast Growth Factor Receptor 4 (FGFR4) (blue).^22^

This initial dataset identified many other protein types proximal to HER2, including ADAM Metallopeptidase Domain 10 (ADAM10), Basal Cell Adhesion Molecule (BCAM), Myelin Protein Zero Like 1 (MPZL1), Lipolysis Stimulated Lipoprotein Receptor (LSR), Sushi Domain Containing 2 (SUSD2), CD36 Molecule (CD36), Protein Butyrophilin Subfamily 3 Member A3 (BTN3A3), and Tyrosine Phosphatase Receptor Type F (PTPRF) (**Figure 1D**). To validate the specificity of HER2-targeted µMap, we performed an *in vitro* fluorescent immunosorbency assay to measure binding of either Tz or an isotype control toward a small panel of immobilized recombinant proteins identified in our dataset (HER2, CD36, INSR, and a negative control using BSA). Only HER2 produced detectable binding in the presence of Tz, strongly indicating that our dataset reflects HER2-specific interactors rather than off-target Tz binding events (**Figure S1**).

Among these identified interactors, ADAM10 has a known direct physical interaction through cleavage of the HER2 ectodomain.^23^ MPZL1 is also known to form a complex with the HER2-associated adaptor GRB2,^24^ while MST1R has been shown to facilitate pathway crosstalk.^25^ In contrast, LSR, CD36, and BTN3A3, although associated within the context of breast cancer biology, have not yet been shown to physically interact with HER2. BCAM and SUSD2 are similarly uncharacterized as HER2 interactors but may modulate HER2 signaling through laminin binding or pathway crosstalk. The identification of PTPRF in proximity to HER2 is particularly interesting because phosphatases typically counteract kinase activity, suggesting a potential regulatory mechanism via removal of inhibitory phosphates. Recent evidence suggests that PTPRF and HER2 may indeed physically interact,^26, 27^ yet the functional relevance of this association is unclear. PTPRF has been shown to act as a tumor suppressor in other malignancies, and its expression has been correlated to RTK dephosphorylation.^28, 29^ Conversely, PTPRF upregulation has been observed in some breast cancer tissues relative to controls, and animal model studies have linked this amplification to increased metastatic potential.^30^

With a robust platform for profiling HER2 associations in place, we next applied µMap across different HER2+ subtypes—including luminal A and B (HR+/HER2+), HER2-enriched (HR-/HER2+), and TNBC (HR-/HER2-)—to explore significant interactome variations. We focused on HER2+ cell lines with comparable receptor expression levels, selecting MDA-MB-361, BT-474, and AU565 as HER2-dependent lines,^31^ and HCC1954 and HCC1419 as models of natural Tz resistance (**Figure 1A**).^32, 33^ Quantitative proteomics of HER2-targeted µMap revealed similar interactors to those identified in AU565 cells while also uncovering distinct trends (**Figures 2A-D**). Setting enrichment thresholds (*p* < 0.05 and Log_2_FC >1.5), we compiled all hits to identify overlap of enriched proteins in each µMap dataset (**Figure 2E**). Notably, a core set of 12 proteins was commonly enriched, potentially indicating the conserved HER2 minimal interactome across cellular contexts. STRING interaction network analysis constructed from these common interactors highlighted functionally related groups, including an RTK centric group, a distinct phosphatase cluster represented by PTPRF and PTPRK, and a cell adhesion group containing BCAM, OCLN, and others (**Figure 2F**). In addition, Gene Ontology (GO) enrichment for these core interactors identified highly relevant pathways to RTK and cell adhesion signaling, reflecting possible conserved functions of HER2 interactions across cancer lines.

**Figure 2.**
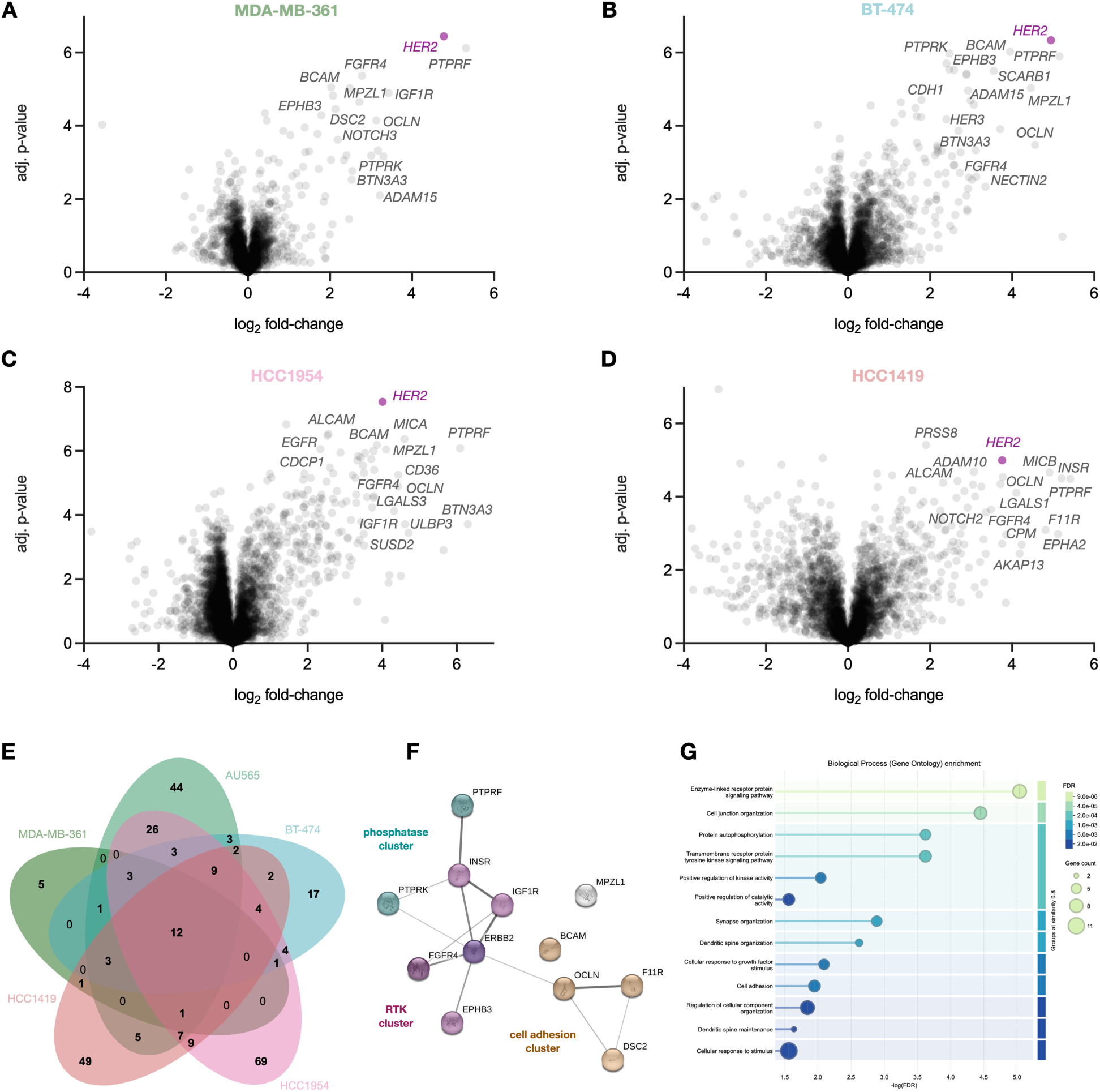
µMap Profiling Identifies Common Features of the HER2+ Interactome. (A-D) Volcano plots displaying µMap proteomics in HER2+ cell lines. (E) Venn diagram of enriched HER2 interactors from each dataset (*p* < 0.05, Log2FC > 1.5). (F) STRING interaction network analysis of the 12 common interactors across all HER2+ cell lines. (G) Biological process (Gene Ontology) enrichment of the common 12 proteins enriched.

### HER2-Galectin Interactions Facilitate Trastuzumab Resistance

We next sought to determine if µMAP could reveal interactome differences between HER2-dependent and independent cell lines. Strikingly, we observed an inverse relationship between HER2 dependency and interactome size, such that an increase in the number of interactors was correlated with a decrease in HER2 dependence (**Figure 3A**). This finding suggests that gain-of-function interactions may underlie Tz-resistance. We initially hypothesized that differential enrichment of RTKs would be readily observable as part of this trend, yet there was no consistent pattern between sensitive or resistant cell lines (**Figure 3B**). In contrast, we observed specific µMap enrichment of several other protein classes, including galectins (LGALS1/GAL1, LGALS3/GAL3), lipid signaling-related proteins (LSR, LDLR, LIPH), and cell adhesion molecules (CDH1, CDH3, CXADR, CELSR2) (**Figure 3C**). Comparison of total protein levels (**Figure 3D**) with HER2 proximity data revealed that for several interactors—including LIPH and LGALS1— higher cellular abundance was correlated with µMap enrichment. However, other proteins showed notable HER2 interaction despite low overall expression, underscoring that while proximity is related to abundance, it is a more nuanced reflection of specific molecular interactions.

**Figure 3.**
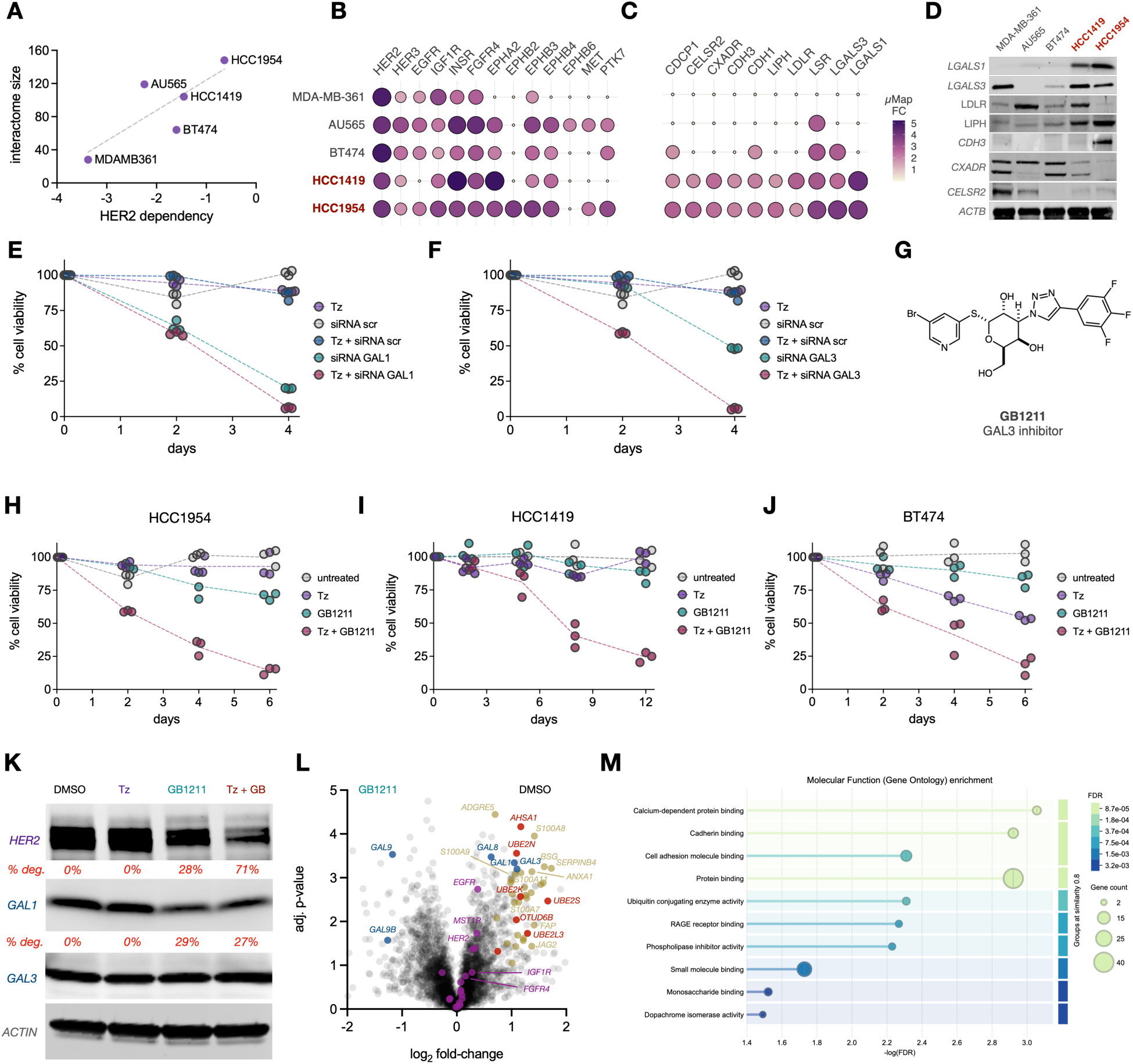
HER2-Galectin Interactions Facilitate Trastuzumab Resistance. (A) DepMap HER2 dependency scores vs measured µMap interactome sizes (*p* < 0.05, Log2FC > 1.5). (B, C) Balloon plot of selected interactors (RTKs (B), other protein types (C)) depicting µMap fold-enrichment change (FC) across all 5 HER2+ cell lines. (D) Western blot analysis of protein expression levels of selected interactors. (E, F) Cell proliferation measurement of HCC1954 cells treated with Tz, GAL1/3 KD, or combinations of each treatment. (G) Chemical structure of GB1211, a small molecule inhibitor of GAL3. (H, I, J) Cell proliferation measurement of HCC1954 (H), HCC1419 (I) and BT-474 cells (J) treated with Tz, GB1211, or in combination. (K) Western blot analysis of HER2, GAL1 and GAL3 expression in treated HCC1954 cells. (L) HER2 µMap proximity labeling of HCC1954 cells treated with GB1211. Galectins (blue), cell adhesion and Ca^2+^ related signaling proteins (beige), RTKs (purple), protein-stability hits (red). (M) Molecular Function enrichment of HER2 interactors displaced upon GB1211 treatment.

We next assessed the functional influence of interactors on cell proliferation and Tz sensitivity. Galectin-1 (GAL1) and Galectin-3 (GAL3) both belong to a larger family of glycan-binding proteins, often upregulated in cancer, that are involved in cell adhesion, signaling, tumor progression, metastasis, and immune escape.^34-37^ GAL3 has also been shown to interact with other RTKs and enhance transduction of the PI3K/AKT signaling pathway.^38^ GAL3 upregulation has also been demonstrated in Tz-resistant HER2+ breast cancer,^39^ yet neither a direct physical interaction between HER2 and galectins nor a mechanistic link to resistant phenotypes has yet been established. Using the Tz-resistant HCC1954 line as a model, we first measured viability over the course of 4 days when treated with either Tz, a non-targeting siRNA control (scrambled, scr), siRNA targeting GAL1 or GAL3, or combinations of each treatment (**Figure 3E, F**). As expected in these resistant cells, Tz treatment alone produced little change in viability, whereas knockdown (KD) of GAL1 or GAL3 alone significantly reduced cell growth over 4 days. Strikingly, combination KD with Tz treatment produced synergistic cytotoxicity that was especially notable upon GAL3 KD (**Figure 3F**).

To further investigate this observation, we explored pharmacological inhibition of galectins as a combination strategy. We identified GB1107 and GB1211 (**Figure S3, 3G**) as potential candidates given their current clinical investigations as small molecule GAL3 inhibitors.^40-42^ We ideally sought compounds that would enhance Tz efficacy while not imparting significant toxicity on their own, so we first evaluated tolerability of both compounds in HCC1954 cells treated with various concentrations for 72 hours. GB1107 displayed noticeable toxicity with increasing doses, while GB1211 was well-tolerated at all concentrations tested and was selected for study (**Figure S3**). We first measured proliferation of HCC1954 cells treated with either Tz (10 µg/mL), GB1211 (10 µg/mL), or a combination of both. Treatment with Tz or GB1211 alone had limited impact on proliferation over 6 days, but significant sensitization and viability decrease were observed in HCC1954 cells treated in combination (**Figure 3H**), supporting the results of genetic knockdown of GAL1/3 and suggesting a galectin-mediated mechanism of Tz resistance. This sensitization and synergistic cytotoxicity effect was confirmed in the Tz-resistant cell line HCC1419 (**Figure 3I**). Surprisingly, we even observed Tz enhancement in sensitive HER2+ lines BT-474 and AU565 (**Figure 3J, S4**). Overall, these findings identify galectins as functional key regulators underlying HER2 dependency and Tz response.

Given these striking results, we next sought to validate the HER2-Galectin interaction and interrogate the mechanism of Tz and GAL3 combination efficacy. We first visually assessed interactions of HER2 with either GAL1 or GAL3 in HCC1954 (high galectin expression) and AU565 (low galectin expression) cells. GAL1 and GAL3 are known to mediate diverse cellular processes and exhibit broad subcellular localization across the nucleus, cytoplasm, cell surface, and extracellular space.^34-37^ Immunofluorescence staining of GAL1/3 and HER2 in HCC1954 cells revealed co-localization with HER2 at the cell membrane and at some locations the cytoplasm (**Figure S5**). In AU565 cells, GAL1 exhibited co-localization with HER2 while GAL3 displayed very faint staining, consistent with our earlier findings on the expression levels of both proteins (**Figures S6, 3D**). To validate physical interactions, we used a proximity ligation assay targeting HER2 and either GAL1 or GAL3. As expected, we observed positive amplification signals for both HER2/GAL3 and HER2/GAL1 in the high-expressing HCC1954 cells compared to negative controls (**Figure S7**). In low-expressing AU565 cells, HER2/GAL1 was detected yet GAL3 signal interactions were not observed, again likely due to significantly lower expression of this galectin (**Figure S8**).

### Galectin Inhibition Disrupts HER2 Stability and Remodels Interactions

Tz resistance can manifest through a variety of mechanisms, including expression of mutated or truncated HER2 isoforms lacking the Tz binding site, or hyper-glycosylation to occlude antibody binding.^3, 43-45^ Given the role of galectins in glycan-binding and the potential effects on HER2, we first assessed direct antibody binding as an initial hypothesis for sensitization. However, in line with our immunofluorescence staining results, Tz binding was unimpeded in HCC1954 cells. Unexpectedly, treatment with GB1211 reduced antibody binding rather than enhancing it, ruling this out as a mechanism for GB1211 synergy (**Figure S9**). We reasoned that HER2 levels may be altered upon GB1211 and/or Tz treatment. Interestingly, GAL inhibition alone resulted in ∼28% degradation of HER2, which increased to over 70% when combined with Tz treatment (**Figure 3K**). GB1211 also led to degradation of GAL1 (∼29%) yet not GAL3 itself, while Tz alone had no impact on the degradation of either HER2 or GAL1/3 in the Tz-resistant HCC1954 cell line. Although GB1211 is selective toward GAL3 (*K*_D_ ∼25 nM), it exhibits binding to other galectins, including GAL1 (*K*_D_ ∼3 µM), in the doses used in our study (∼19 µM) (**Figure S10**).^41^ This may explain Tz enhancement in cells expressing lower amounts of GAL3 (BT-474 and AU565). GB1211 may also trigger a unique destabilizing conformational change or disrupt interactions critical for the integrity of GAL1—events that may not occur with GAL3. Overall, the substantially enhanced and synergistic degradation of HER2 observed with the combination of GB1211-mediated GAL3 inhibition and Tz treatment is strongly implicated as a major mechanistic driver for the efficacy of this co-treatment strategy.

We next probed interactome-level changes to HER2 upon galectin inhibition with GB1211. HCC1954 cells treated with either DMSO or GB1211 for 72 hours were harvested, and then µMap proximity labeling of HER2 was performed on both samples followed by quantitative proteomic analysis (**Figure 3L, Table S6**). As expected, GAL3 itself was displaced from the HER2 interactome, providing primary validation that the inhibitor functions as intended. Interestingly, GAL1 and GAL8 were also displaced, again suggesting some promiscuity in GB1211 across galectins. GAL9 was also surprisingly HER2-enriched only when GB1211 was present, possibly stemming from reduced competition for shared or similar glycosylated binding sites on or near HER2, or the exposure of new interaction surfaces within the altered HER2 complex. RTK heterodimer enrichment was relatively constant in both conditions, reflecting our earlier µMap data across different HER2+ cancers (**Figure 3B**). GO enrichment of HER2 interactors displaced upon GAL inhibition (log_2_FC > 1, *p* < 0.05) revealed significant functional associations with cell adhesion, migration, glycolysis, calcium-binding, as well as ubiquitin-related pathways (**Figure 3M**), providing clues into how GAL3 inhibition might enhance Tz and promote HER2 degradation. In particular, there was consistent displacement of several annexin isoforms (ANXA1, ANXA3, ANXA5) as well as Profilin-1 (PFN1), Basigin (BSG), Coactosin-like protein 1 (COTL1), Jagged Canonical Notch Ligand 2 (JAG2) and Migration and invasion enhancer 1 (MIEN1), among others, strongly suggesting that galectins provide important links between HER2 and specific sets of interactors.^46, 47^ We also observed significant displacement of S100 calcium-binding proteins, suggesting disruption of HER2’s ability to leverage pro-tumorigenic and glycolytic signaling, with additional effects on adhesion and metabolic dynamics relevant to cancer progression.^48, 49^ In line with our results showing enhanced HER2 degradation with galectin inhibition, we observed displacement of ubiquitin-conjugating enzymes UBE2K, UBE2L3, UBE2N and UBE2S, deubiquitinase OTUD6B, as well as co-chaperone AHSA1. These UBE isoforms are all implicated in non-degradative K63-linked ubiquitination,^50, 51^ strongly suggesting that GAL3 facilitates these events to stabilize HER2 or promote its signaling functions. AHSA1 is a co-chaperone that stimulates HSP90,^52^ which in turn stabilizes HER2.^53, 54^ Modest displacement of the main HSP90 proteins themselves (HSP90AA1, HSP90AB1) was also observed (**Table S6**), further supporting the idea that galectins mediate or stabilize HER2-chaperone interactions and providing a rationale for the observed HER2 degradation. Overall, our interactomic analysis reveals that galectin inhibition severely impairs HER2 by simultaneously displacing adhesion and pro-tumorigenic signaling networks while critically disrupting protective ubiquitination signatures and chaperones—ultimately leading to HER2 destabilization.

### Galectin Inhibition with Trastuzumab Induces Broad Cellular Reprogramming

In parallel, we evaluated global proteomic changes in HCC1954 cells treated with GB1211, Tz, or in combination (**Tables S7-S10**). Using identical thresholds for up- and downregulated protein expression (log_2_FC > 0.5, *p* < 0.05), we compared global expression level changes across each treatment. Although Tz-treatment alone exhibited little change in expression patterns (**Figure S11, Table S7**), GB1211-mediated galectin inhibition, both alone and in combination with Tz, resulted in significant global changes. We first analyzed the effects of GB1211 alone (**Figure 4A**, **Table S8**). We noted downregulation of a large number of genes regulating cell cycle, proliferation, and DNA metabolism, including mitosis regulation (CCNB2, CDK1, NDC80, NUF2, KIF4A, KIF2A, SPC24/25),^55^ DNA replication (DNMT1, POLE, POLA2, RRM2, TYMS, UHRF1, HELLS),^56^ and DNA repair (RECQL5, RAD54B/L2, FANCA, UBE2T).^57^ We did not observe significant cytotoxicity in cells treated with GB1211 alone (**Figure 3H-J**), suggesting that its activity does not result in severe or complete arrest but may sensitize cells to Tz by compromising their division capabilities. Consistent with our western blot analysis (**Figure 3K**), HER2 was downregulated with GB1211 treatment, along with several other RTKs found in the dataset (**Figure 4A**), suggesting a common galectin-mediated scaffolding mechanism that may stabilize these receptors. Key signaling mediators STAT3, JAK2, MAPK8 and Rho GTPase regulators (ARHGEF40, ARHGAP8/18/29) were also suppressed.^58, 59^ Cell adhesion and cytoskeletal proteins were also downregulated (AHNAK, CLDN8, DSP, SDC1, DSC2, NECTIN1, SMTN), supporting a trend in which galectin inhibition alters cell architecture with reduced adhesion.^60, 61^

**Figure 4.**
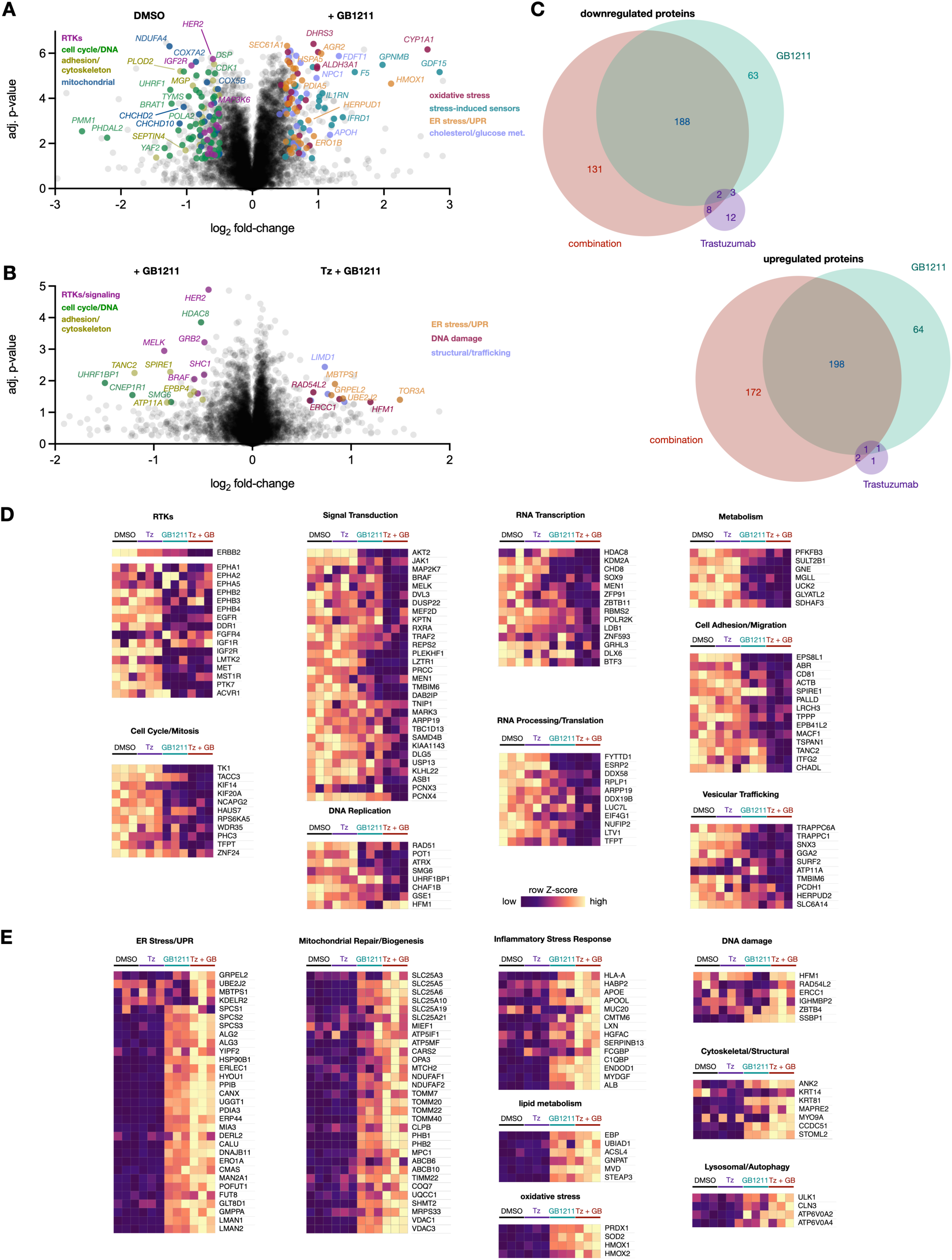
Global Proteomic Changes under Galectin Inhibition and Trastuzumab. (A, B) Volcano plots displaying quantitative global proteomic analyses of cells treated with either DMSO or GB1211 (A) or GB1211 vs combination treatment (B). Protein types are colored according to the legend. (C) Venn diagram of down- and up-regulated protein expression levels across Tz, GB1211, or combination treatment. (D, E) Heatmap of quantitative proteomics (row Z score) across treatments showing (D) downregulated and (E) upregulated expression across gene categories.

We also identified genes that were upregulated in response to galectin inhibition (**Figure 4A, Table S8**). We observed significant enrichment of proteins involved with endoplasmic reticulum (ER) stress and Unfolded Protein Response (UPR), including chaperones and folding enzymes such as HSPA5, DNAJC3, DNAJC10, PDIA4/5/6, ERO1B, and HERPUD1.^62, 63^ Inhibition of galectin function likely disrupts protein folding and ER homeostasis due to altered glycoprotein processing or accumulation of misfolded proteins. Oxidative stress response genes were also increased, including CYP1A, HMOX1, SQSTM1, SOD2, CAT, GCLC and GCLM,^64, 65^ likely a consequence of ER stress and metabolic shifts. Upregulation of SQSTM1, along with WIPI1, LAMP1, LAMP2, and CTSL (Cathepsin L), suggests autophagy activation and enhanced lysosomal activity,^66, 67^ potentially to clear damaged organelles or aggregated proteins. Interestingly, we also observed upregulation of the cholesterol biosynthesis pathway, including FDFT1, SQLE, LSS, DHCR7, DHCR24, NSDHL, and EBP. This suggests a major remodeling of the membrane lipid composition, perhaps to cope with membrane stress and altered receptor trafficking.^68-70^ Glucose metabolic shifts were also apparent, including increased expression of PDK4, ALDH3A1/2, ALDH3B1 and DHRS3/4/9, indicating increased metabolic and detoxification processes.^71^ Lastly, upregulation of stress-response modulators, including GDF15, IGFBP3, IDO1, and NDRG1, among others,^72^ was also observed, likely in response to cellular damage. Overall, this analysis revealed that galectin inhibition orchestrates profound cellular reprogramming of proliferation, energy production, metabolic adaptation, proteostasis, and stress response pathways. Critically, it also destabilizes and reduces the expression of HER2 along with several other RTKs.

We directly compared global expression between cells treated with GB1211 alone versus in combination with Tz (**Figure 4B, Table S10**). In line with our earlier results showing Tz-induced HER2 degradation (**Figure 3K**), addition of Tz resulted in further downregulation of HER2 and several associated signaling partners (GRB2, SHC1).^24, 73^ Interestingly, key oncogenic kinases BRAF and MELK were also decreased, indicating suppression of MAPK/ERK proliferation pathways. Lowered expression of cell adhesion and membrane organization markers (TSPAN1, EPB41 TANC2, ITFG2, SPIRE1)^74, 75^ suggests further structural destabilization along with potentially increased phosphatidylserine exposure due to ATP11A suppression,^76^ likely rendering cells more susceptible to apoptosis and clearance. Downregulation of DNA replication factors (UHRF1BP1, SMG6 and HDAC8)^77, 78^ along with concomitant upregulation of DNA damage repair proteins (HFM1, RAD54L2, ERCC1 and IGHMBP2)^79^ provides evidence that Tz combination treatment inflicts higher levels of DNA damage and replication stress than GB1211 alone. HER2 signaling is known to support DNA repair and cell survival and may contribute to the therapeutic effect of Tz.^80, 81^ Thus, by promoting HER2 degradation, galectin inhibition may sensitize cells to damage or reduce repair capacity. Intensified proteotoxic burden and ER dysfunction was again evident with Tz addition, with further upregulation of UBE2J2,^82^ MBTPS1, the Hsp70 co-chaperone GRPEL2, and TOR3A (**Figure 4B**).^83^ As noted above, GB1211 alone does induce ER stress (**Figure 4A**); the addition of Tz and resulting HER2 degradation likely increases the accumulation of misfolded or aberrant proteins, accelerating ER-associated degradation (ERAD) and enhancing UPR toxicity.

As expected, when compiling all differentially expressed proteins from Tz, GB1211, or combination treatment relative to DMSO, we observed the most similarity among the galectin-inhibited conditions (**Figure 4C**). We then isolated differential expression signatures unique to the combination treatment and categorized them for both down- (**Figure 4D**) and up-regulated (**Figure 4E**) genes across each treatment. The synergistic efficacy of galectin inhibition combined with Tz reflects a multi-faceted response, characterized primarily by suppression of HER2 along with several RTKs at enhanced levels compared to either agent alone. Additionally, further downregulation of critical signaling pathways involving AKT2, JAK1, BRAF, and MELK is evidence for diminishing proliferation and survival. Concurrently, the combination uniquely disrupts genomic maintenance and cell cycle capacities by further suppressing essential genes in DNA replication and mitosis, while also impairing crucial cell adhesion/migration and vesicular trafficking machinery (**Figure 4D**). While the galectin inhibitor alone initiates cellular stress, the combination treatment intensifies upregulation of genes indicative of severe ER stress/UPR, profound mitochondrial dysregulation and biogenesis attempts, and a heightened DNA damage response (**Figure 4E**). Overall, the accelerated HER2 degradation alongside accumulation of cellular damage, pathway disruption, and compromised repair and survival pathways likely explains the observed synergistic effect.

### HER2 Interacts with PTPRF and is Upregulated Across Breast Cancers

After mapping HER2+ interactomes and identifying dependency mechanisms, we next applied µMap across a broader panel of breast cancer subtypes—including the HR-/HER2+ lines MDA-MB-453 and JIMT-1; the luminal B (HR+/PR+/HER2+) line ZR-75-1; the luminal A (HR+/HER2-) line MCF-7; and the TNBC model MDA-MB-231—as well as the normal mammary cell line MCF10A (**Figure 5A-F, S12, Tables S12-17**). In HER2+ cell lines, µMap revealed similar interactors to those identified in our earlier experiments (**Figure 1D, 2**)—including Ephrin receptor variants, RTKs IGF1R and INSR, ADAM proteases, BCAM, and MPZL1—reinforcing their roles as core members of the HER2 proximal interactome. Similarly, prominent enrichment of protein tyrosine phosphatase F (PTPRF) was again observed in the HER2+ lines MDA-MB-453, JIMT-1, and ZR-75-1 (**Figure 5A,B, S12**). This trend was especially notable as we profiled HER2-cell lines MCF-7 and MDA-MB-231 (**Figure 5C,D**). We also profiled the normal MCF10A mammary epithelial cell line (**Figure 5E**) and PTPRF was not detected, suggesting a cancer-specific association with HER2. Combined comparisons with our earlier datasets to reveal that HER2 interactome sizes (*p* < 0.05, Log_2_FC > 1.5) scale with expression but also correlate with genetic dependency in HER2+ cell lines (**Figure 5F**).

**Figure 5.**
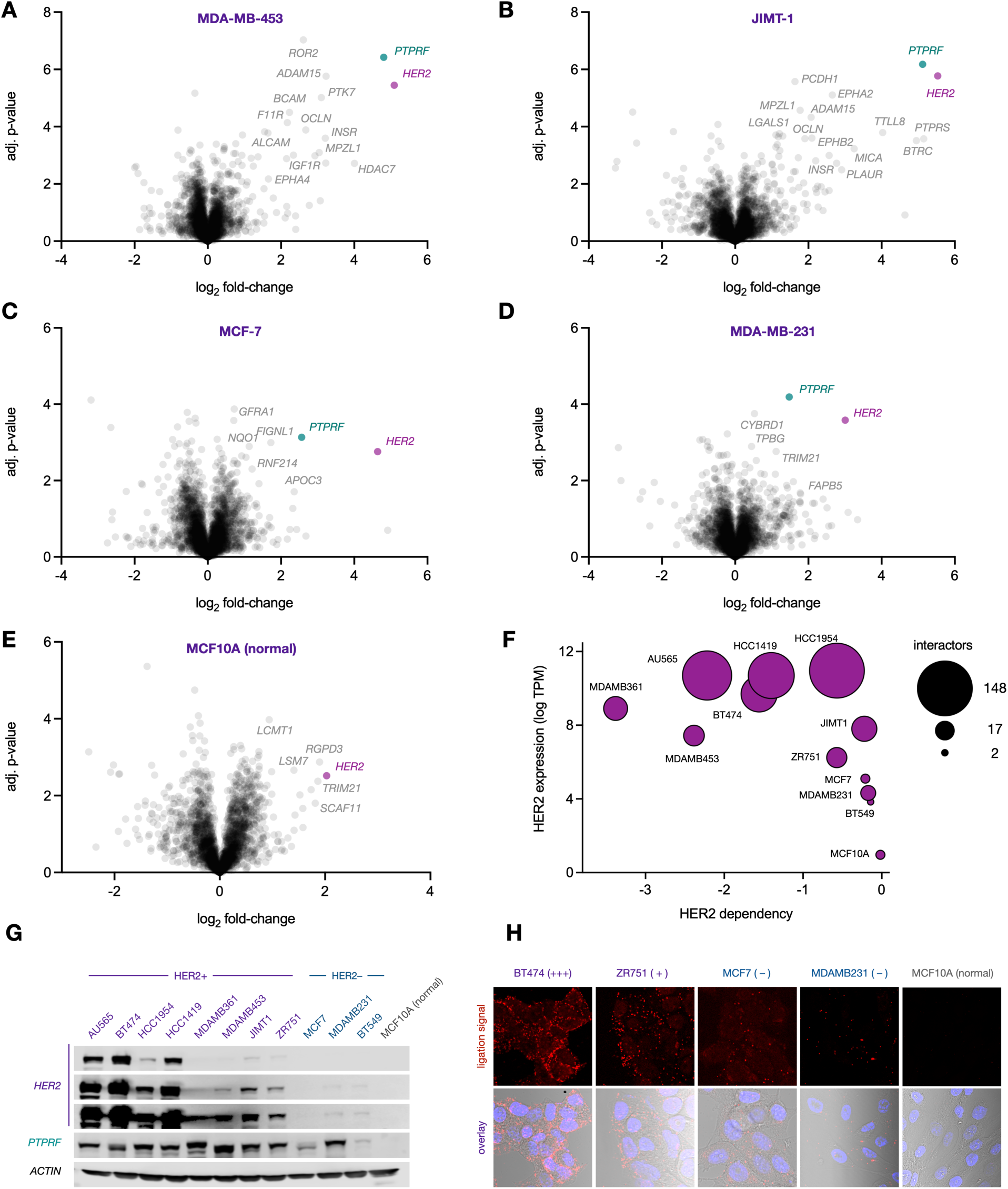
PTPRF Interacts with HER2 and is Upregulated Across Cancers. (A-E) Volcano plots displaying quantitative proteomics analysis of HER2 µMap in MDA-MB-453, JIMT-1, MCF-7, MDA-MB-231 and MCF10A cell lines. (F) DepMap HER2 dependency scores vs measured µMap interactome sizes (*p* < 0.05, Log2FC > 1.5). (G) Western blot analysis of HER2 and PTPRF expression levels in selected breast cancer cell lines. (H) HER2 and PTPRF proximity ligation assay in selected breast cancer cell lines.

To further explore the interaction between HER2 and PTPRF, we first measured expression across our panel of cell lines (**Figure 5G**). As expected, HER2 expression was correlated with HER2+ cells, with lower expression in HER2-cell lines and undetectable levels in the normal MCF10A line. Intriguingly, PTPRF was robustly expressed across all cancer cell lines irrespective of HER2 status, and again was undetectable in normal MCF10A cells. These data strongly suggest PTPRF is upregulated during carcinogenesis and implies a broader functional role for PTPRF in breast cancer biology. We also validated the *in situ* physical interaction between HER2 and PTPRF using a proximity ligation assay, observing abundant red ligation signals at the cell membrane in HER2+ BT-474 cells, with progressively fewer signals in ZR-75-1, MCF-7, and MDA-MB-231 cancer cells (**Figure 5H, S13**). Importantly, the interaction was again undetectable in normal MCF10A cells, correlating with our µMap results and observed protein expression levels (**Figure 5E, G**), reinforcing that the HER2-PTPRF interaction is a significant feature in cancer cells, even those with low overall HER2 levels.

### PTPRF Co-Expresses with HER2 and is Amplified Across Cancer Types

Given this consistent relationship between HER2 and PTPRF, we next examined expression correlations across larger datasets. Previous studies have observed cancer-specific upregulation of PTPRF^84, 85^ as well as positive correlations with HER2 expression.^86, 87^ We first plotted HER2 against PTPRF expression from all breast cancer lines in DepMap^18^ and observed a moderate but biologically significant correlation (*R*^2^ = 0.25, **Figure S14**). While not as robust compared to oncogenes with known co-expression, including GRB7^88^ (*R*^2^ = 0.79) or STARD3^89^ (*R*^2^ = 0.59, **Figures S15,16**), this correlation was comparable to the heterodimer HER3 (*R*^2^ = 0.34, **Figure S17**, and was in fact much more significant than either EGFR (*R*^2^ = 0.03,) or IGF1R (*R*^2^ = 0.01, **Figures S18, 19**).

To gain a broader understanding of PTPRF and HER2 co-expression, we analyzed a large scRNA-seq dataset profiling normal breast tissue along with all major cancer subtypes.^90^ As expected, visually plotting HER2 expression, we saw upregulation enriched in HER2+ subtypes, with minimal signal across ER+, PR+, and TNBC samples (**Figure 6A**). Interestingly, PTPRF displayed more widespread expression, with strong signal overlap in HER2+ samples along with significant signals in ER+ and TNBC (**Figure 6B**), consistent with the observed upregulation of PTPRF across breast cancer lines. Mean scRNA-seq expression levels further support cancer-associated PTPRF upregulation, with ∼2-3-fold increases observed across subtypes compared to normal cells. In contrast, HER2 amplification was mainly observed in HER2+ tumors (**Figure 6C**). Expression correlation between the two genes across all cell types also indicated a positive but modest overlap (*ρ* 0.20 Jaccard index “J” 0.30) (**Figure 6D**). Examining all candidate genes across cell types, PTPRF was among the five genes most highly co-expressed with HER2, displaying similarity indices equivalent to HER2-HER3 and higher than GRB7^88^ or MIEN1^91^ (**Figure S20**). Notable co-occurrence and similarity were observed within the HER2+ subtype (*ρ* 0.45, J 0.53), while normal breast tissue samples displayed much weaker linkage and overlap (*ρ* 0.12 J 0.20), suggesting limited co-occurrence in normal breast tissue cells. Collectively, these data support cancer-associated upregulation and co-expression between PTPRF with HER2. Although the genetic mechanisms for this co-expression are not fully clear at present and could reflect complex regulatory feedback loops, this relationship underscores a potential functional importance in HER2-related signaling and breast cancer oncogenesis.

**Figure 6.**
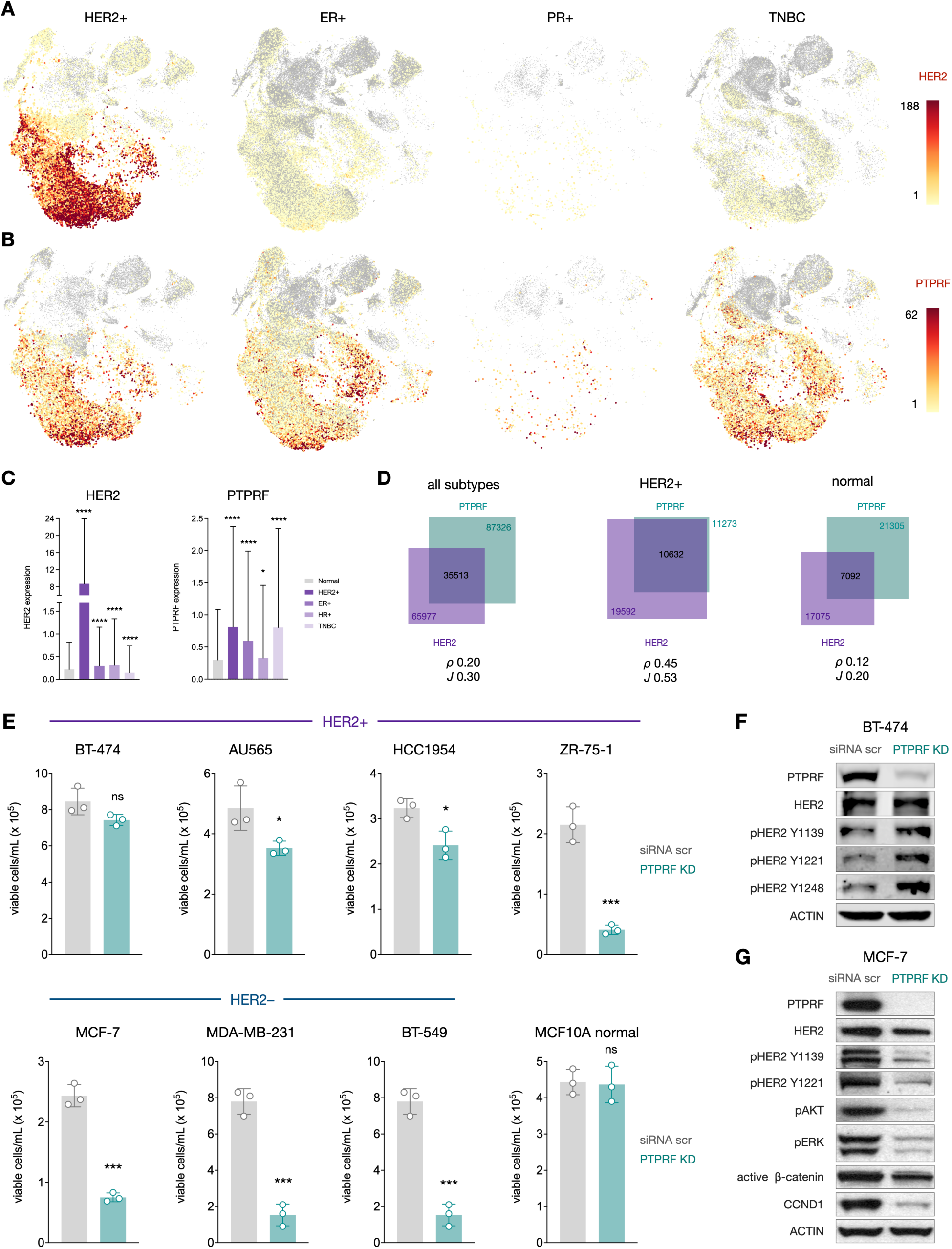
PTPRF is upregulated and co-expressed with HER2 and functionally impacts cell signaling and proliferation. (A, B) scRNA-seq expression of HER2 and PTPRF across cancer types. (C) Mean expression of HER2 and PTPRF between breast and cancerous subtypes. Error bars denote S.D. (*p* values denote t-test vs normal tissue (* < 0.05, **** < 0.0001)). (D) Co-expression similarity between HER2 and PTPRF across subtypes. *ρ* values denote linear expression correlation and J values display overlap similarity in positive expressing cells. (E) PTPRF siRNA KD and cell viability measurement across cancer and normal cells. (F, G) Western blot analysis of BT-474 and MCF-7 cells after PTPRF KD.

### PTPRF Silencing Influences Cell Proliferation and HER2 Phosphorylation

To elucidate the functional roles of PTPRF in breast cancer, we performed siRNA-mediated KD of PTPRF across multiple cancer cell lines, along with the normal MCF10A line (**Figure S21**). Intriguingly, the impact of PTPRF silencing on cell viability varied significantly depending on cellular context (**Figure 6E**). HER2– lines exhibited the most reduction in viability upon PTPRF KD, with ER+ MCF-7 and TNBC lines MDA-MB-231 and BT-549 showing significant decreases. In contrast, responses among HER2+ cell lines were more heterogeneous: ZR-75-1 cells experienced dramatic viability loss comparable to HER2– lines, whereas AU565 and HCC1954 cells exhibited statistically significant yet minor decreases in viability. Notably, BT-474 cells were unaffected by PTPRF KD (**Figure 6E**). These differences may reflect the high oncogenic output and increased number of HER2 interactors in the HER2-amplified setting, potentially providing a degree of redundancy to PTPRF-supported pathways. Conversely, the sensitivity of ZR-75-1 cells suggests that PTPRF can still play an important role, even in the context of HER2 amplification, perhaps by regulating distinct survival pathways. The moderate sensitivity of AU565 and HCC1954 cells could indicate an intermediate level of dependency, wherein both HER2 signaling and PTPRF-supported pathways contribute to cell viability.

Importantly, PTPRF KD did not significantly affect MCF10A normal-like epithelial cells, consistent with our earlier findings that PTPRF expression is undetectable in this line (**Figure 5G**). The differential sensitivity to PTPRF loss across cancer cell lines suggests that while PTPRF is broadly expressed, its functional necessity becomes heightened in malignant contexts, potentially as cancer cells become rewired toward specific PTPRF-supported signaling axes. The pronounced selectivity over normal cells, combined with the impact on TNBC and ER+ cell lines—which often lack effective targeted therapies or exhibit resistance–highlights the broader therapeutic potential of PTPRF inhibition.

To investigate the molecular mechanisms underlying PTPRF function and cell viability, we performed western blot analysis on BT-474 and MCF-7 cells following PTPRF KD. In BT-474 cells, PTPRF silencing led to a notable increase in HER2 phosphorylation at signaling residues Y1139, Y1221, and Y1248^92^ (**Figure 6F**). In contrast, MCF-7 cells showed reduced phosphorylation at Y1139 and Y1221, along with a mild reduction in total HER2 levels (**Figure 6G**). However, it is worth noting that both cell lines experienced significant decreases in proliferation and overall confluency compared to control KD. Consistent with the observed reduction in cell viability in PTPRF KD cells, we observed profound downregulation of key proliferative signaling pathways—in particular, suppression of PI3K/AKT and MAPK/ERK as well as decreased levels of active β-catenin and Cyclin D1 (**Figure 6G**). These results strongly suggest that in MCF-7 cells, PTPRF is crucial for maintaining the activity of these fundamental signaling networks that drive cell survival, proliferation, and cell cycle progression. While some studies have characterized PTPRF as a tumor suppressor in certain malignancies,^39^ more recent findings reveal an oncogenic function in certain cancers where it actively promotes proliferation and survival by sustaining MAPK/ERK and Wnt/β-catenin pathways.^30, 93, 94^

The differential effects of PTPRF KD in BT-474 versus MCF-7 cells underscore the context-dependent and complex roles of this enzyme, which may act as a phosphatase for HER2, potentially fine-tuning its activation state. The lack of a strong viability effect in BT-474 cells suggests that these cells can tolerate fluctuations in phosphorylation or that the role of PTPRF is more nuanced than simple growth suppression. Conversely, in MCF-7 cells, PTPRF is important for sustaining critical pro-survival pathways. Evidence suggests that PTPRF associates directly with β-catenin and cadherins or may act upstream of the destruction complex,^93, 94^ and the precise targets and mechanisms by which PTPRF supports these pathways is a key area for further investigation. The dramatic reduction in viability in MCF-7 cells upon PTPRF KD is likely a direct consequence of the collapse of these PTPRF-dependent signaling networks. Overall, these data highlight PTPRF as a complex regulator of cell signaling and a potential therapeutic target in ER+ and TNBC tumors, where it appears to sustain oncogenic signaling independent of high HER2 expression.

## DISCUSSION

HER2 and other RTKs orchestrate intricate interaction networks that are central to breast cancer progression, yet the mechanisms underpinning HER2 dependency and therapeutic resistance remain opaque.^1, 2, 95^ Similarly, the role of these networks in HER2– tumors is not well understood.^3^ Here, we leverage the high-resolution capabilities of µMap to comprehensively chart the HER2 interactome across 11 breast cancer cell lines spanning all major subtypes. Our findings not only provide an extensive interactomic resource but also unveil novel mechanisms of intrinsic Tz resistance and identify galectin family proteins and the protein tyrosine phosphatase PTPRF as novel therapeutic targets.

In Tz-resistant HER2+ models, galectins were identified as uniquely enriched HER2 interactors. We demonstrated that both genetic and pharmacological inhibition of galectins, particularly GAL3, can restore Tz sensitivity in resistant cell lines and enhance efficacy in sensitive lines. Mechanistically, this is driven by destabilization and enhanced degradation of HER2, dissociation of adhesion machinery and protective chaperone systems, and initiation of ER and mitochondrial stress pathways. These findings underscore a key role for galectins in HER2 signaling and oncogenesis, warranting further clinical investigation of galectin inhibitors as a combination strategy to overcome Tz resistance.

Our pan-cancer µMap profiling platform also repeatedly identified PTPRF as a HER2 interactor with upregulation detected in cancer cell lines over normal tissue. Functionally, PTPRF silencing profoundly suppressed proliferation in ER+ and TNBC cell lines, and this effect correlated with downregulation of key pro-survival pathways, including PI3K/AKT, MAPK/ERK, and Wnt/β-catenin. These results suggest that PTPRF is important for maintaining these fundamental oncogenic signaling networks. While PTPRF knockdown did increase phosphorylation in HER2-driven BT-474 cells, the lack of a viability effect in this line alongside the profound dependency in HER2-negative lines points to complex, context-dependent functions. Future studies should therefore focus on precisely defining how PTPRF may influence HER2 phosphorylation states and understanding context-dependent therapeutic strategies for exploiting this interaction. Concurrently, identifying the scope of PTPRF interactors and substrates is critical to mechanistically understanding its role in cell proliferation and signaling regulation, particularly in HER2– subtypes where its KD had a profound impact.

In conclusion, this work leverages µMap proximity labeling to deliver an unprecedented insight into the HER2 interactome across the breast cancer landscape. These findings significantly advance our understanding of HER2 signaling complexity in normal and diseased tissue. The extensive interactomic data provided herein serves as a valuable resource for the community, paving the way for further mechanistic discoveries and the development of innovative cancer therapies.

## LIMITATIONS OF THE STUDY

While this work provides insights into the HER2 interactome and identifies novel therapeutic avenues, certain limitations should be acknowledged. Firstly, our HER2 interactome profiling, although utilizing the high-resolution µMap platform, does capture proximity rather than direct and/or functional binding. However, the small labeling radius of the technique allows capture of only the most proximal interactors, whereas conventional immunoprecipitation-based approaches can be misleading by virtue of isolating large protein complexes—thus attributing indirect interactions as proximal and direct. Future studies employing complementary biochemical validation and structural analyses of interactors throughout the dataset will be important to confirm all direct interactions and elucidate their functional consequences.

Secondly, while our investigation spanned 11 breast cancer cell lines across various subtypes, this represents a fraction of the extensive heterogeneity inherent in human breast cancer. These *in vitro* 2D culture models may also not fully recapitulate the complex tumor microenvironment and signaling dynamics of *in vivo* disease. Future work should involve expanding µMap profiling to a broader array of patient-derived xenografts, organoids, or tissue slides to validate and explore these findings. Moreover, the identification of galectins and tyrosine phosphatases as therapeutically actionable targets in breast cancer warrants their investigation as potential regulators in other models of resistance or additional cancer types. While we identified PTPRF as a cancer-specific HER2 interactor and demonstrated its role in cancer cell viability, the precise molecular mechanisms governing this action require more in-depth investigation. Elucidating its specific substrates and upstream regulators in different contexts will be crucial for fully understanding its therapeutic potential across the diverse landscape of breast cancer and potentially other malignancies. Deeper mechanistic understanding of PTPRF’s pro-oncogenic functions could also facilitate development of potent and selective small molecule phosphatase inhibitors or targeted protein degradation modalities to effectively neutralize its contributions to cancer cell survival and proliferation.

## Supporting information

Experimentals and Supplemental Figures

## Resource availability

### Lead contact

Data are available upon request to lead contact, David W. C. MacMillan.

## Acknowledgments

Research reported in this work was supported by the Princeton Branch of the Ludwig Institute for Cancer Research, National Institute of General Medical Sciences of the National Institutes of Health (R35GM134897), the Princeton Catalysis Initiative, and kind gifts from Merck, Pfizer, Janssen, Bristol Myers Squibb, Genentech, and Genmab. S.D.K. acknowledges the NIH for postdoctoral fellowships (1F32GM142206 and 1K99GM154140). J.G. thanks the Deutsche Akademie der Naturforscher Leopoldina (LPDS 2023-03) for financial support. We would also like to thank Gary S. Laevsky and Sha Wang of the Confocal Imaging Facility, a Nikon Center of Excellence, in the Department of Molecular Biology at Princeton University for instrument use and technical advice. Illustrations were created with BioRender.com.

## Author contributions

S.D.K., J.A.B., J.D.R., and D.W.C.M. conceived the study; J.D.R., and D.W.C.M supervised the study; S.D.K, J.A.B., and D.C.M. performed most µMap and LC-MS/MS analysis, cell viability, and western blotting experiments; S.D.K. performed immunofluorescence and proximity ligation assays; J.G. synthesized and characterized GB1211. S.D.K., J.A.B., J.D.R., and D.W.C.M. wrote the manuscript with comments from all the authors.

