## Supplementary material for "Micromapping (µMap) of HER2 Across Human Breast Cancers: Photocatalytic Proximity Labeling Identifies Primary Resistance Mechanisms and Functional Interactors": Experimentals and Supplemental Figures

#### Supporting Information

**Steve D. Knutson**,[1,2†] **Jacob A. Boyer**,[2,3,4†] Danielle C. Morgan,[1,2] Johannes Großkopf,[1,2] Joshua D. Rabinowitz,[2,3,4] and David W. C. MacMillan[1,2*]

[1]Merck Center for Catalysis at Princeton University, Princeton, New Jersey, 08544, USA
[2]Department of Chemistry, Princeton University, Princeton, New Jersey, 08544, USA
[3]Lewis-Sigler Institute for Integrative Genomics, Princeton University, Princeton, New Jersey 08544, USA
[4]Ludwig Institute for Cancer Research, Princeton University, Princeton, New Jersey 08544, USA

† equal contributions

*Corresponding author

#### Supporting Figures

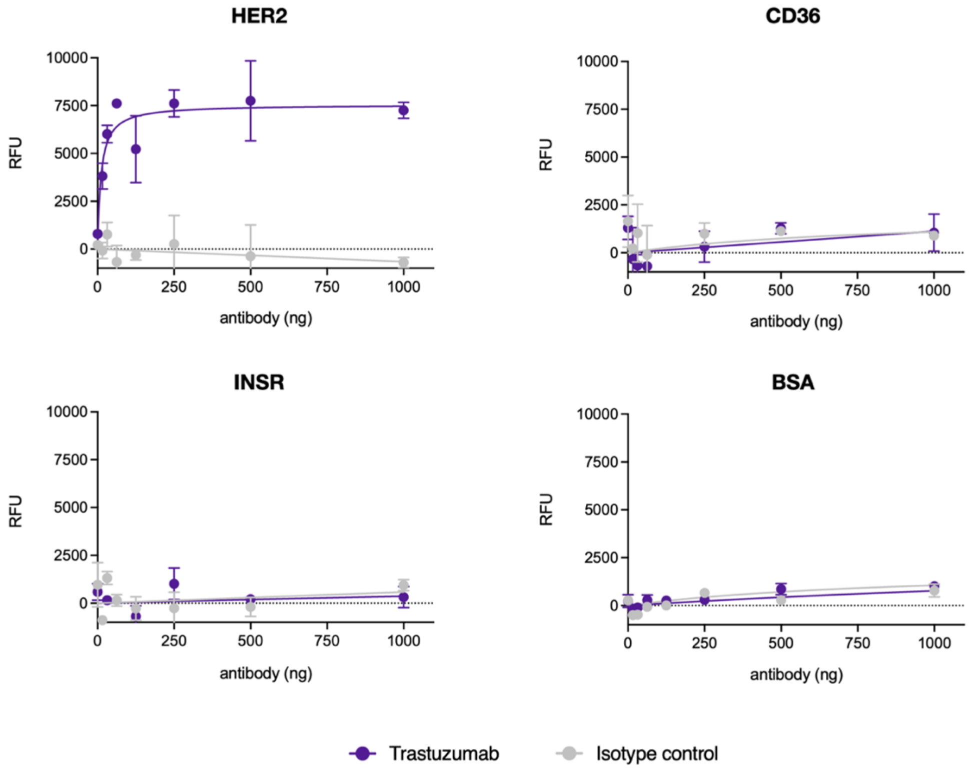

**Figure S1:** *In vitro* trastuzumab binding assay.

Binding of either trastuzumab (purple) or human IgG isotype control (gray) with immobilized recombinant proteins. Values represent net relative fluorescence units (RFU) of each well (n = 2) as measured with a platereader (BioTek). Error bars denote standard deviation (S. D.). Line of best fit was generated in GraphPad Prism.

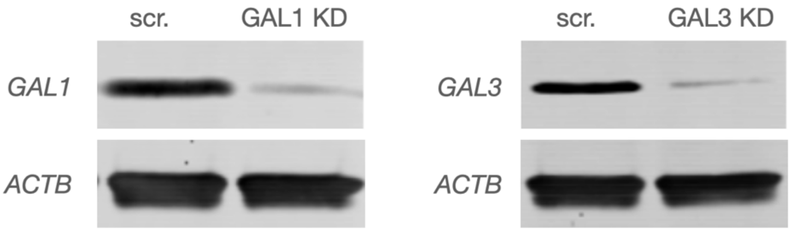

**Figure S2:** Validation of GAL1 and GAL3 knockdown with siRNA

Western blot analysis of whole cell lysates from HCC1954 cells treated with either a non-targeting scrambled siRNA or targeting GAL1 or GAL3.

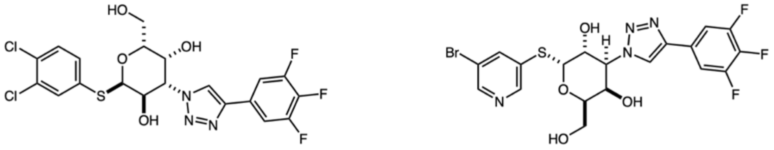

GB1107 GB1211

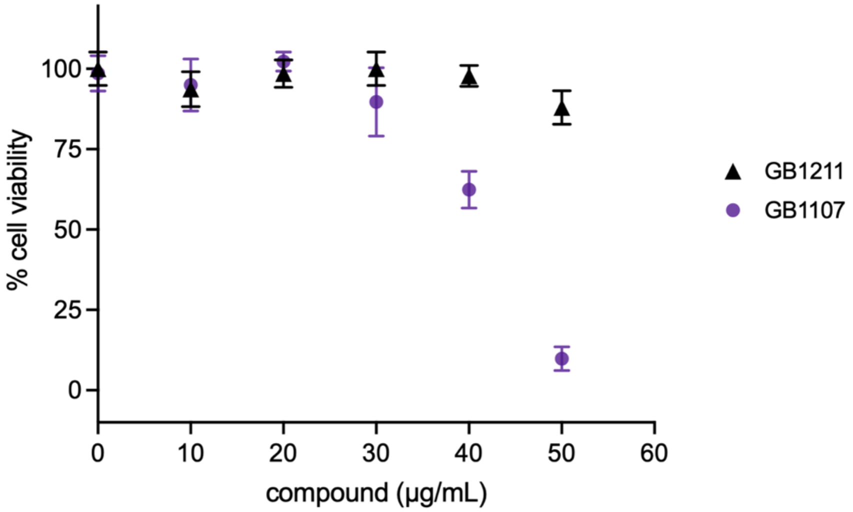

**Figure S3:** Cell proliferation measurement of HCC1954 cells treated with galectin inhibitors.

Chemical structures of GB1107 and GB1211. Cell viability measurement of cells treated with indicated concentrations of either compound. Values represent viability compared to vehicle (DMSO) wells (n = 3). Error bars denote standard deviation (S. D.).

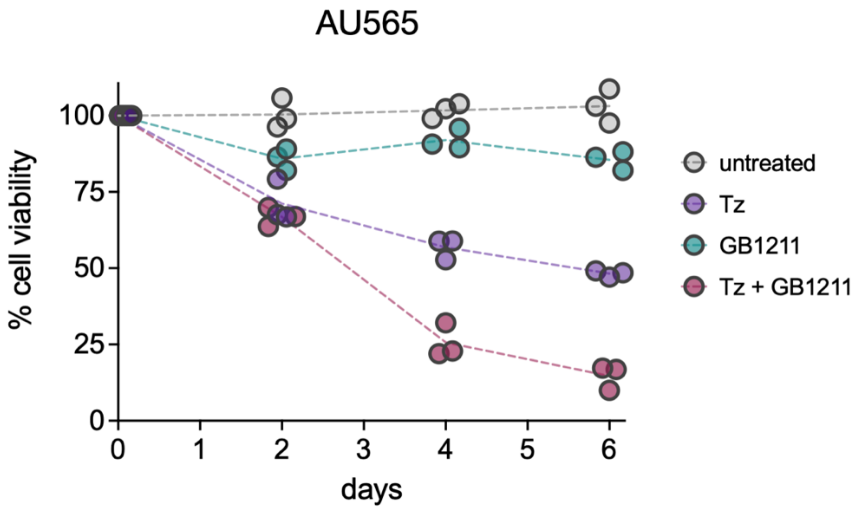

**Figure S4:** Cell proliferation measurement of galectin inhibition in AU565 cells

Cell viability measurement of AU565 cells when treated with galectin inhibitors with and without trastuzumab. Values represent mean viability compared to vehicle (DMSO) wells from three separate experiments (n = 9 total wells).

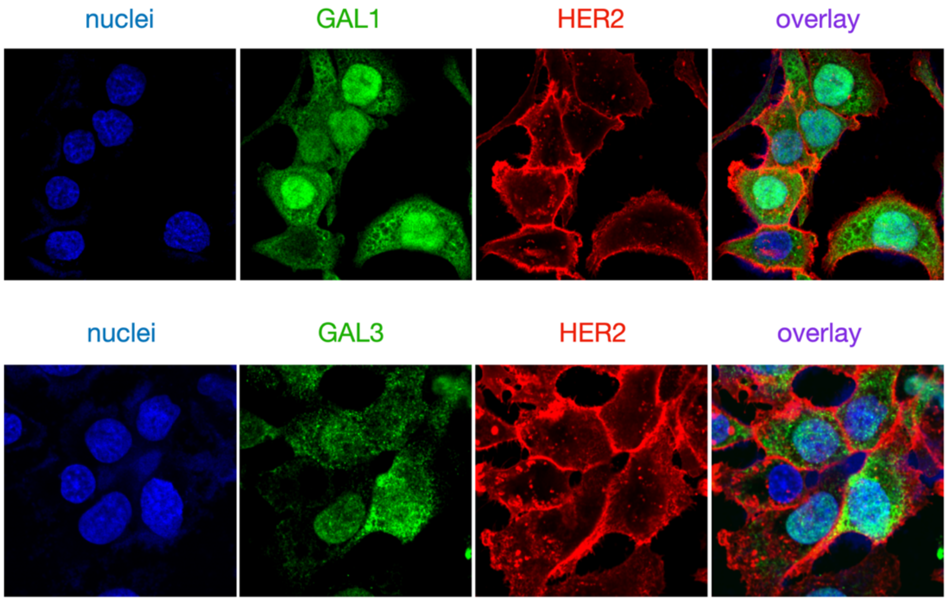

**Figure S5:** HER2 and galectin immunofluorescence in HCC1954 cells.

Representative immunofluorescence images of subcellular localization of GAL1/3 (green) and HER2 (red). Nuclei are stained with DAPI (blue).

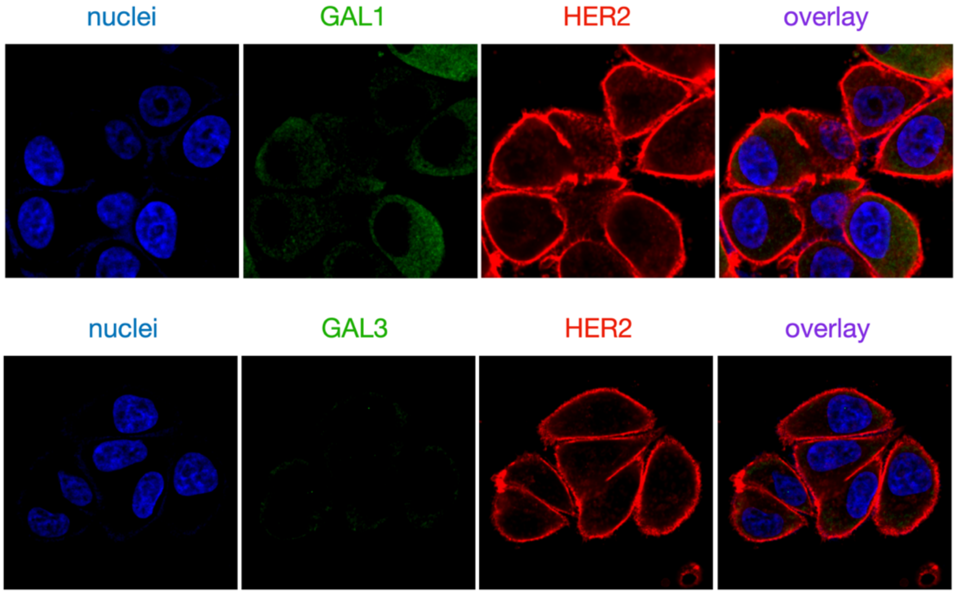

**Figure S6:** HER2 and galectin immunofluorescence in AU565 cells.

Representative immunofluorescence images of subcellular localization of GAL1/3 (green) and HER2 (red). Nuclei are stained with DAPI (blue).

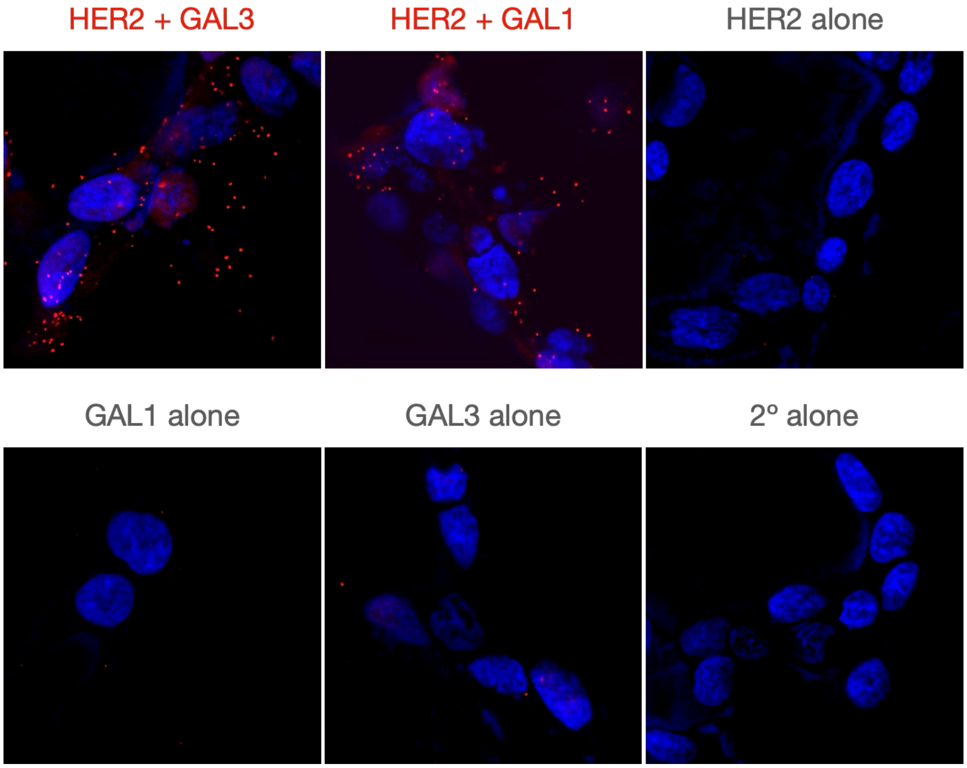

**Figure S7:** HER2 and galectin proximity ligation in HCC1954 cells.
PLA signals are shown in red and nuclei in blue.

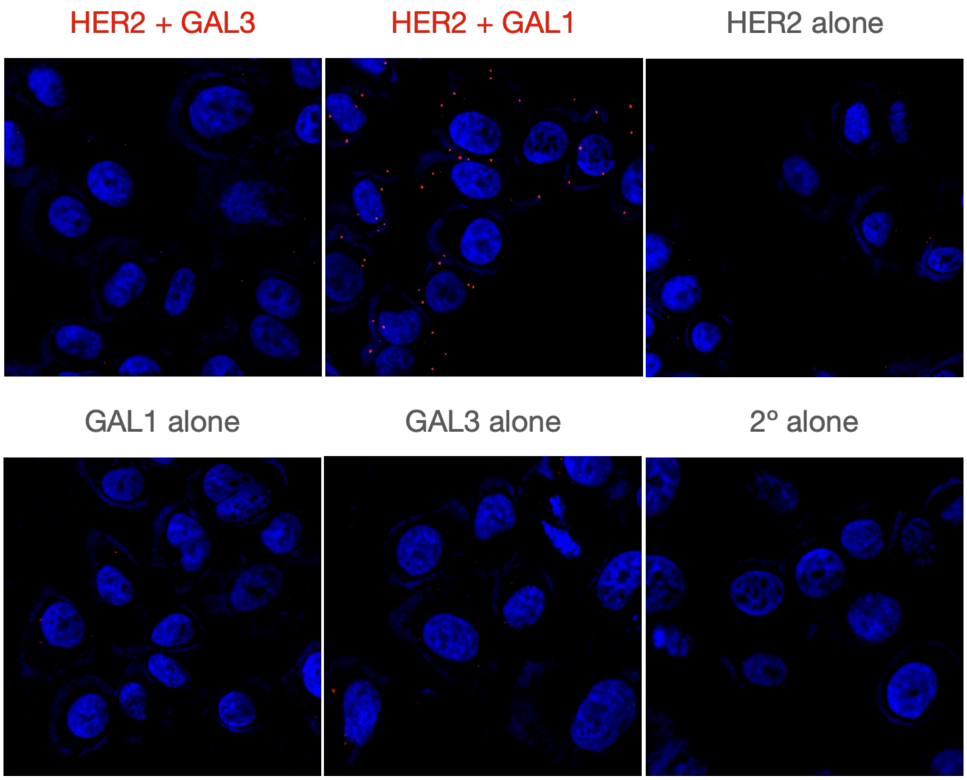

**Figure S8:** HER2 and galectin proximity ligation in AU565 cells.
PLA signals are shown in red and nuclei in blue.

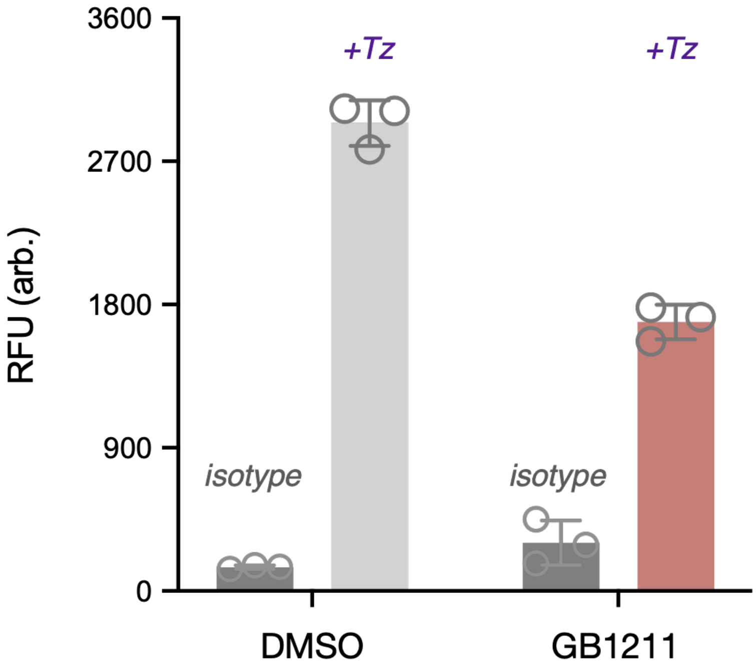

**Figure S9:** Trastuzumab cell binding assay with galectin inhibition.

HCC1954 cells treated with either DMSO or GB1211 for 4 days were incubated with either an human isotype control or trastuzumab followed by a fluorescent secondary antibody to measure binding. Values represent mean from three technical replicates from three separate binding reactions. Values represent net relative fluorescence units (RFU) of each well as measured with a platereader (BioTek). Error bars denote standard deviation (S. D.).

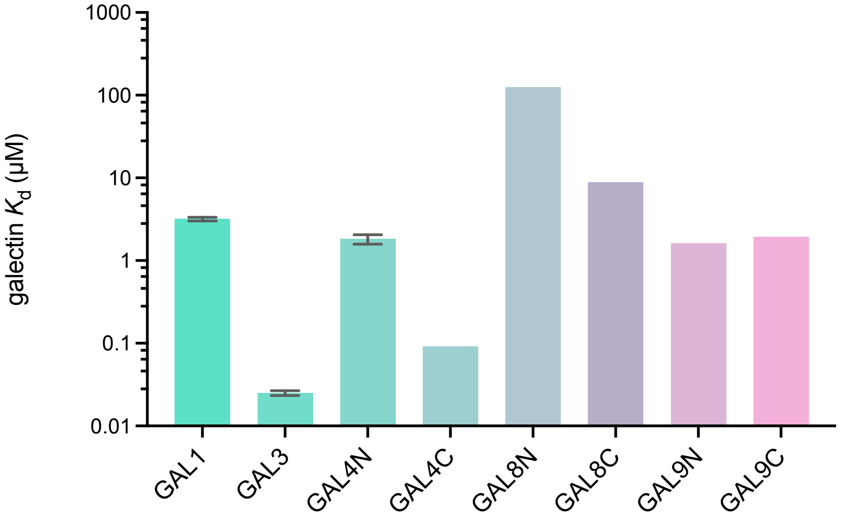

**Figure S10:** Binding affinities of GB1211 for different galectin isoforms.

*In vitro* binding affinities as measured by fluorescence polarization with recombinant proteins, adapted from data in *Journal of Medicinal Chemistry*, **2022**, *65*(19), pp.12626-12638. Error bars denote S.D.

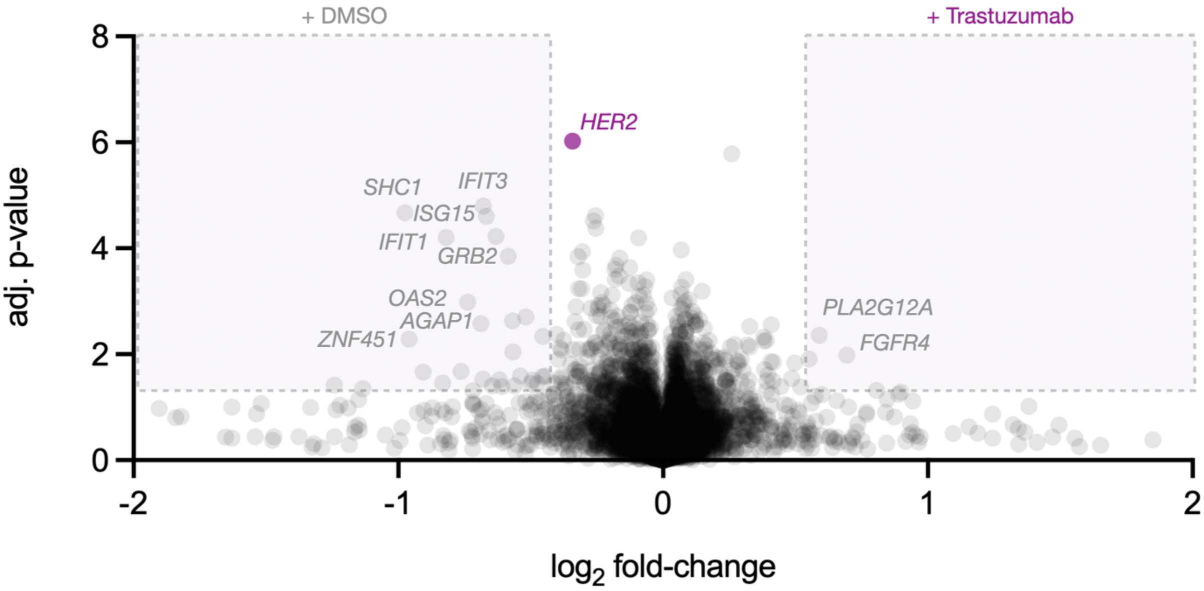

**Figure S11:** Global proteomics of HCC1954 cells treated with trastuzumab.

Volcano plot of Quantitative LC/MS proteomics data of HCC1954 whole cell lysate treated with either DMSO or trastuzumab.

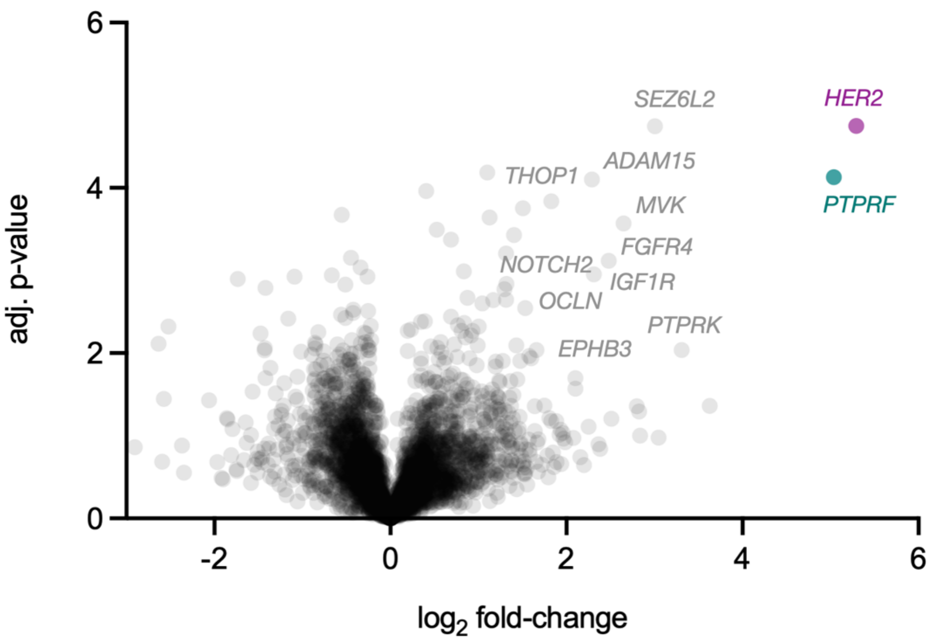

**Figure S12:** HER2 μMap proximity labeling in ZR-75-1 cells.

Volcano plot of Quantitative LC/MS proteomics data of HER2-targeted μMap.

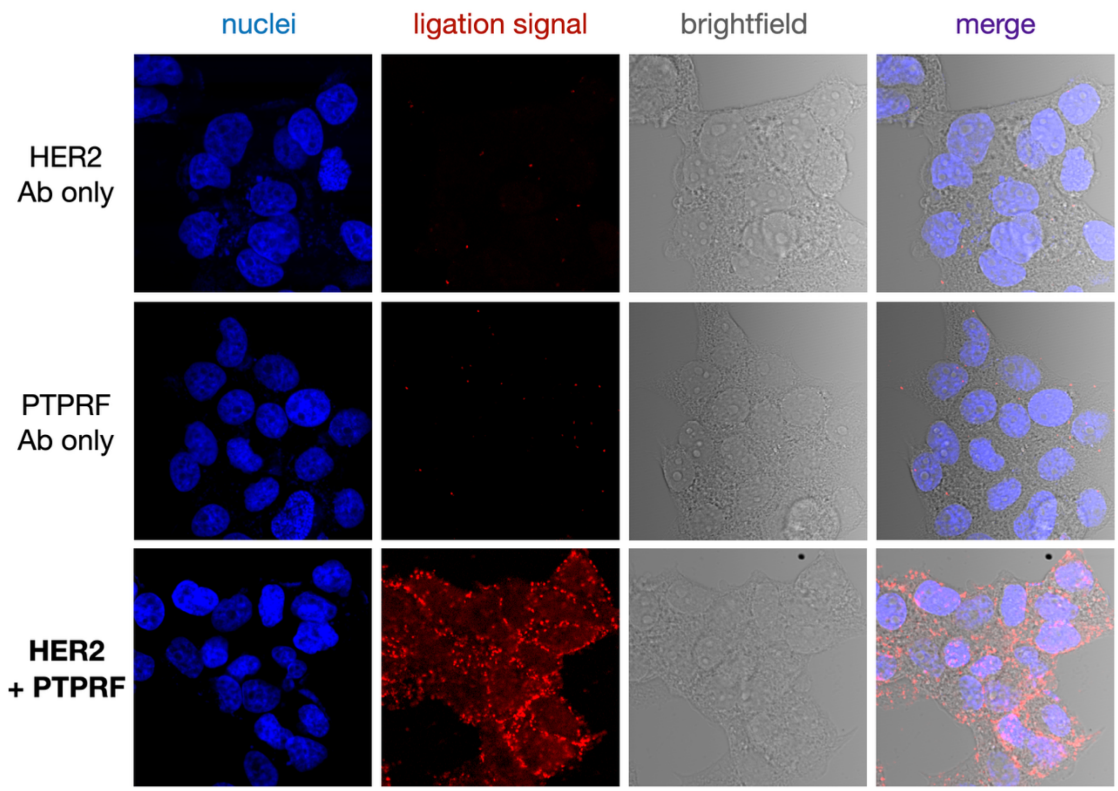

**Figure S13:** HER2 and PTPRF proximity ligation validation in BT474 cells.
PLA signals are shown in red and nuclei in blue.

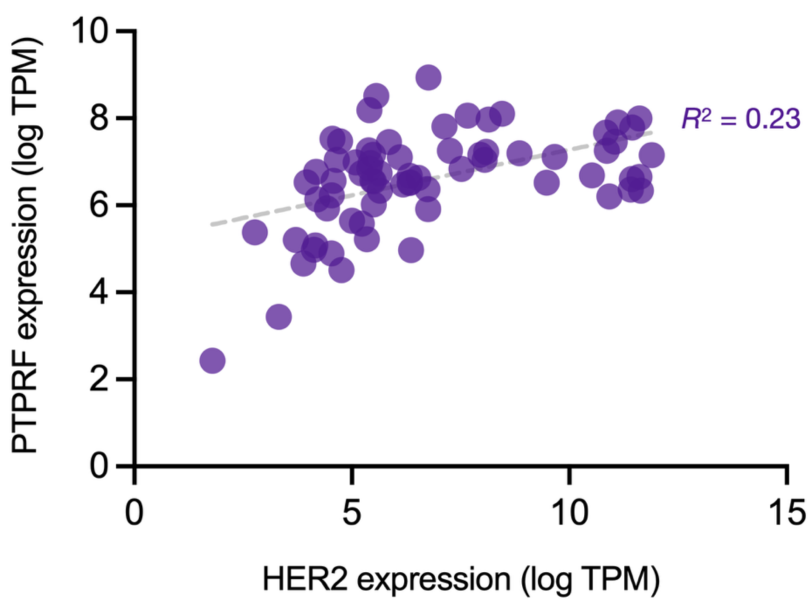

**Figure S14:** HER2 and PTPRF expression across breast cancer cell lines.

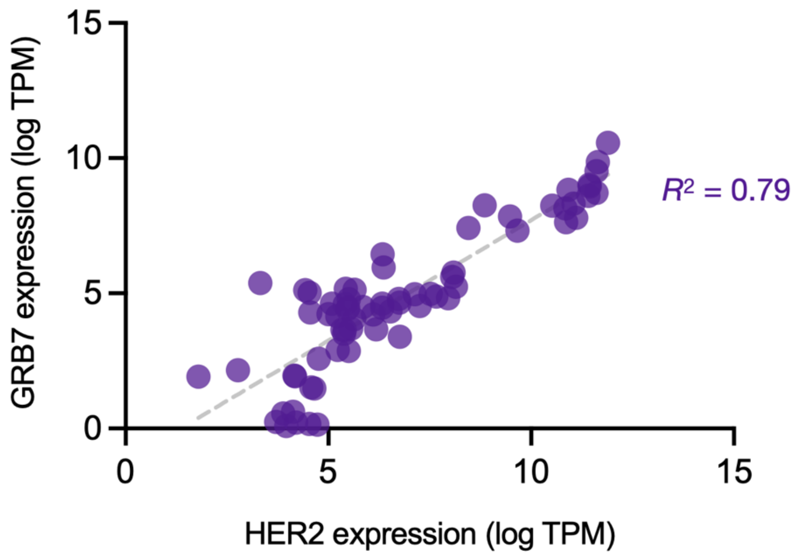

**Figure S15:** HER2 and GRB7 expression across breast cancer cell lines.

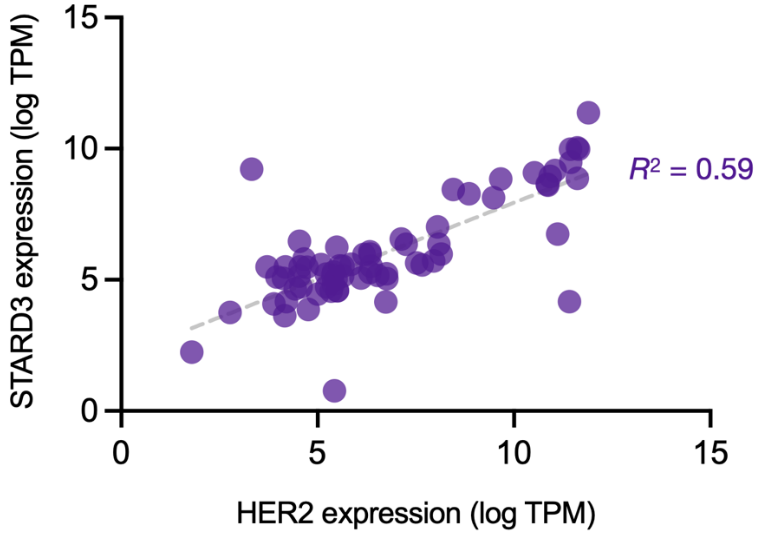

**Figure S16:** HER2 and STARD3 expression across breast cancer cell lines.

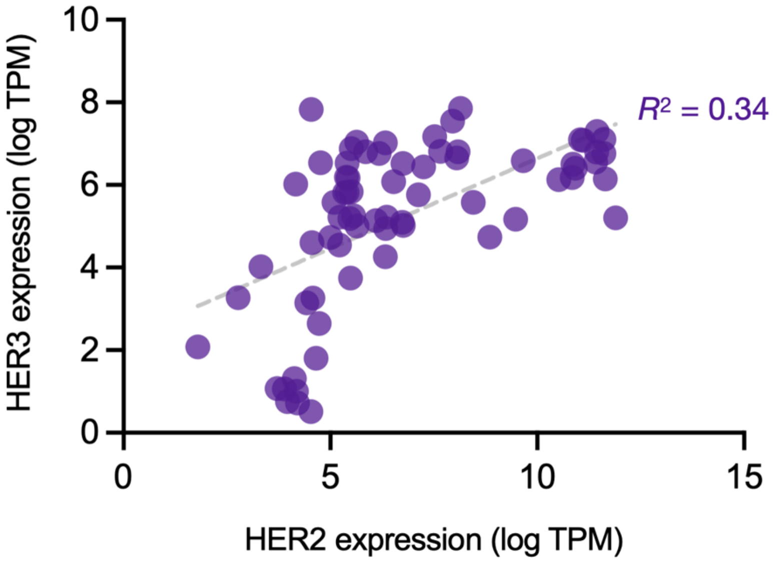

**Figure S17:** HER2 and HER3 expression across breast cancer cell lines.

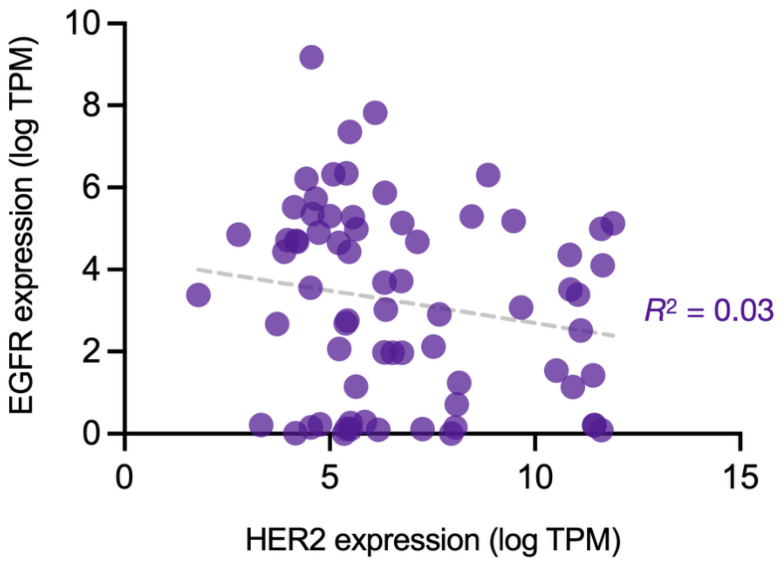

**Figure S18:** HER2 and EGFR expression across breast cancer cell lines.

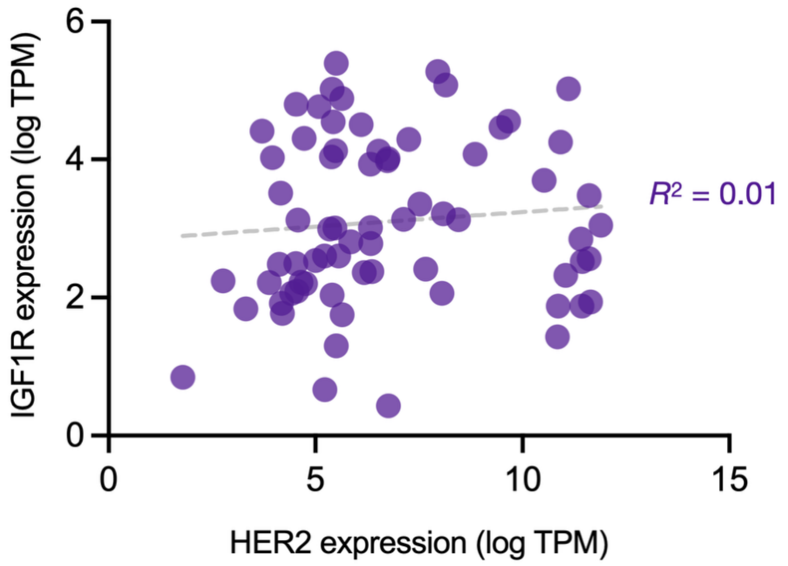

**Figure S19:** HER2 and IGF1R expression across breast cancer cell lines.

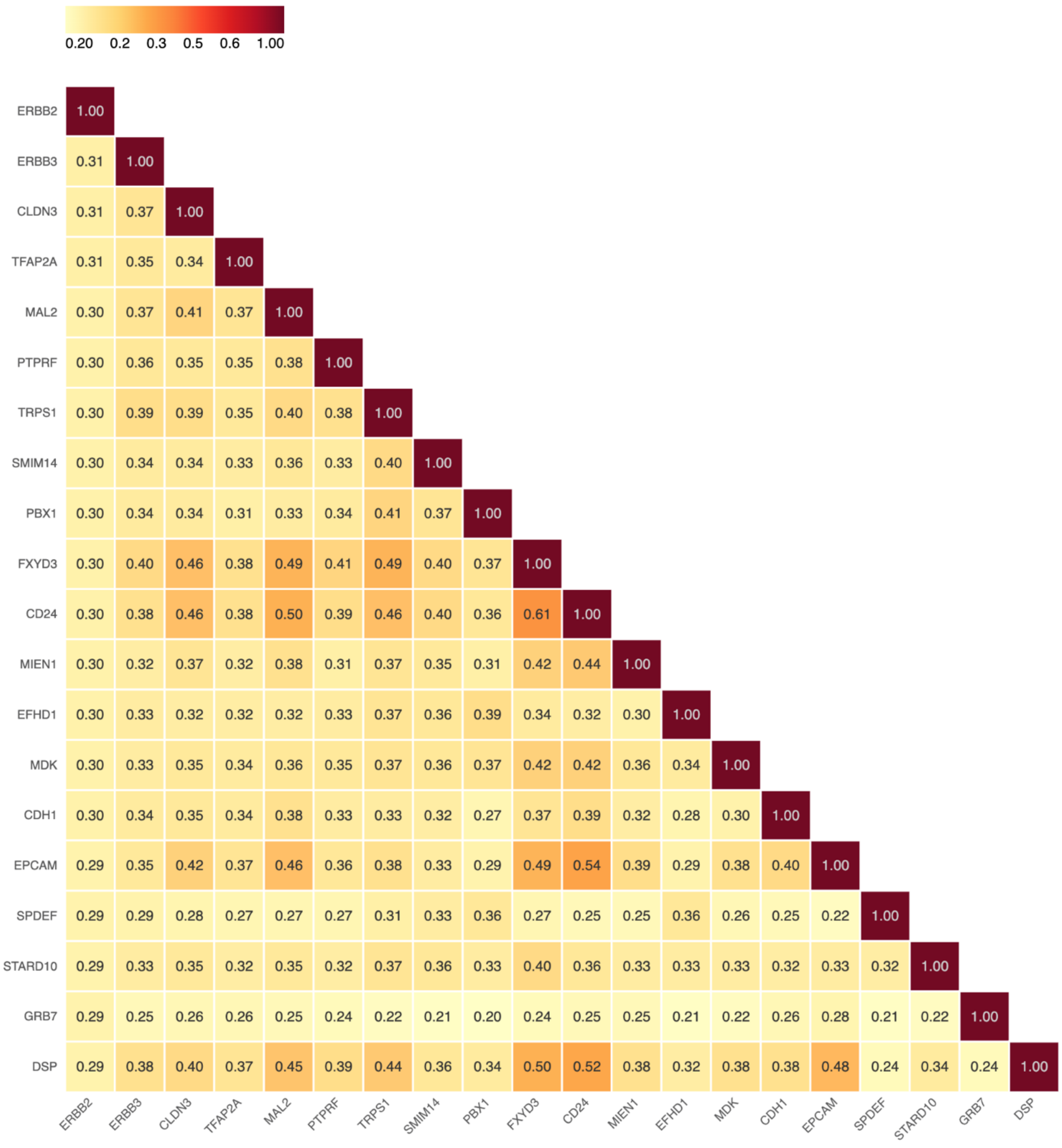

**Figure S20:** Top 20 co-expressed genes with HER2 across all breast cancers and normal tissue. Individual squares denote Jaccard (J) similarity scores.

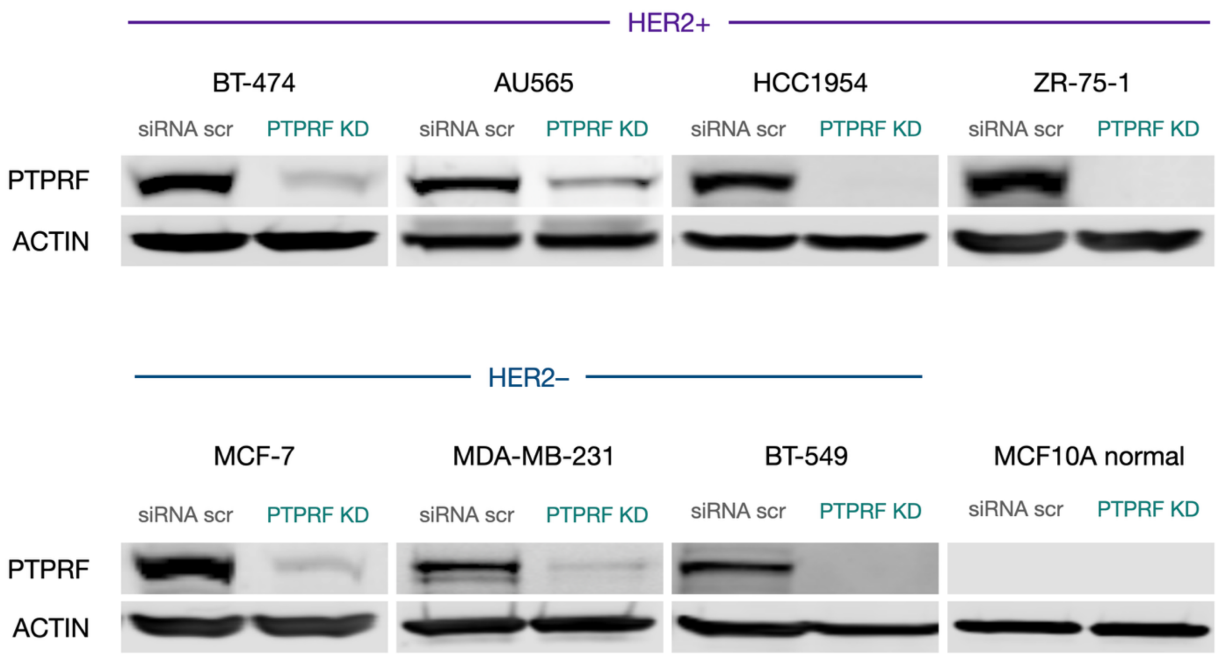

**Figure S21:** Western blot analysis of PTPRF knockdown across cancer cell lines.

### Methods

#### Key Resources Table

| Reagent or Resource | Source | Identifier |
|---|---|---|
| **Cell lines** | | |
| AU565 | ATCC | CRL-2351 |
| BT474 | ATCC | HTB-20 |
| MDA-MB-361 | ATCC | HTB-27 |
| MDA-MB-453 | ATCC | HTB-131 |
| HCC1419 | ATCC | CRL-2326 |
| HCC1954 | ATCC | CRL-2338 |
| JIMT1 | AddexBio | C0006005 |
| ZR-75-1 | ATCC | CRL-1500 |
| MCF-7 | ATCC | HTB-22 |
| MDA-MB-231 | ATCC | HTB-26 |

| | | |
|---|---|---|
| BT549 | ATCC | HTB-122 |
| MCF 10A | ATCC | CRL-10317 |
| **Cell culture reagents** | | |
| Penicillin-Streptomycin | Gibco | 15140122 |
| Fetal Bovine Serum | Gibco | A5670701 |
| DMEM, high glucose | Gibco | 11965084 |
| RPMI 1640 Medium, HEPES | Gibco | 22400089 |
| MEGM® Mammary Epithelial Cell Growth Medium BulletKit | Lonza | CC-3150 |
| Leibovitz's L-15 Medium | Gibco | 11415114 |
| TrypLE™ Express Enzyme (1X), no phenol red | Gibco | 12604013 |
| **Proximity labeling reagents** | | |
| Human IgG Isotype Control | Thermo Fisher | 31154 |
| Human anti ErbB2 Clone 4D5-8 (Trastuzumab Biosimilar) | Bio-Rad | MCA6092 |
| Dulbecco's PBS (1X DPBS) | Thermo Fisher | 14190094 |
| Amicon 10 kDa ultrafiltration concentrator (0.5 mL) | MIllipore | UFC5010 |
| BupH Carbonate-Bicarbonate 0.2 M pH 9.2 | Thermo Fisher | 28382 |
| Azido-PEG12-NHS ester | BroadPharm | BP-21607 |
| DBCO-Ir photocatalyst | This paper | DOI:10.1021/jacs.2c11094 |
| Zeba™ Spin Desalting Columns and Plates, 40K MWCO | Thermo Fisher | A57761 |
| Pierce™ 660nm Protein Assay Reagent | Thermo Fisher | 22660 |
| Bovine Serum Albumin | Sigma | A7030 |
| Biotin-PEG3-diazirine | This paper | DOI:10.1126/science.aay4106 |
| Blue light photoreactor | Acceled | Photoreactor m2 |
| Streptavidin, Alexa Fluor™ 555 Conjugate | Themo Fisher | S32355 |
| Black 96 well plates | Thermo Fisher | 237108 |
| Mem-PER™ Plus Membrane Protein Extraction Kit | Thermo Fisher | 89842 |
| Protease Inhibitor Cocktail | Sigmas | 11873580001 |
| RIPA Lysis Buffer 10 X | Sigma | 20-188 |
| Pierce™ BCA Protein Assay | Thermo Fisher | 23225 |
| Pierce™ Streptavidin Magnetic Beads | Themo Fisher | 88817 |
| **Proteomics Reagents** | | |
| Sodium Dodecyl Sulfate (SDS) | Invitrogen | 15525017 |
| Ethanol, 200 proof | Decon Labs | 2701 |
| Sodium Chloride | Sigma | S9888 |
| Urea | Sigma | 51456 |
| Thiourea | Sigma | T7875 |
| Biotin, free acid | Sigma | B4639 |
| Ammonium Bicarbonate | Sigma | A6141 |
| DTT | VWR | M109 |
| Iodoacetamide | Ambeed | 144-48-9 |

| | | |
|---|---|---|
| MS-grade trypsin | Thermo Fisher | 90057 |
| Formic acid | Fisher Scientific | A117-50 |
| Axygen 1.7 mL Microcentrifuge Tubes | Corning | MCT-175-L-C |
| 0.22-micron spin filter column | CoStar | 98231-UT-1 |
| HPLC-grade water | Fisher Scientific | W64 |
| LC-MS grade Acetonitrile | Thermo Fisher | 85188 |
| Sera-Mag™ Carboxylate SpeedBeads | Cytiva | 45152105050250 |
| Sera-Mag™ Carboxylate SpeedBeads | Cytiva | 65152105050250 |
| HEPES Bioultra grade | Sigma | 54457 |
| Nonidet™ P40 Substitute solution | Sigma | 98379 |
| Sodium deoxycholate | Sigma | 264103 |
| Glycerol | Fisher Scientific | G33-500 |
| EDTA (0.5 M), pH 8.0 | Invitrogen | AM9260G |
| **Western Blotting** | | |
| Criterion TGX tris-glycine polyacrylamide gel cassettes 4-20% | Bio-Rad | 5671093 |
| Nitrocellulose membranes (0.45 μm) | Bio-Rad | 1620115 |
| 4X Laemmli loading buffer | Bio-Rad | 1610747 |
| 2-Mercaptoethanol | Gibco | 21985023 |
| iBright™ Prestained Protein Ladder | Thermo Fisher | LC5615 |
| Tris base | MP Biomedicals | 11TRIS01KG |
| Glycine | Chem-Impex | 00163 |
| **Antibodies** | | |
| HER2/ErbB2 Antibody | CST | 2242 |
| phospho HER2 (Y1139) Antibody | Abcam | ab53290 |
| phospho HER2 (Y1221/1222) Antibody | CST | 2243 |
| phospho HER2 (Y1248) Antibody | CST | 2247 |
| Actin Antibody | Thermo Fisher | MA1-744 |
| LGALS1 Antibody | CST | 12936 |
| LGALS3 Antibody | CST | 87985 |
| LDLR Antibody | ProteinTech | 66414-1-Ig |
| LIPH Antibody | ProteinTech | 16602-1-AP |
| CDH3 Antibody | CST | 2189 |
| CXADR Antibody | Novus | NBP1-88192 |
| CELSR2 Antibody | CST | 47061T |
| GRAMD4 Antibody | ProteinTech | 24299-1-AP |
| PTPRF Antibody | CST | 61611 |
| phosphoAKT Antibody | CST | 13038 |
| phosphoERK Antibody | CST | 4370 |
| Active Beta Catenin Antibody | CST | 8480 |

| | | |
|---|---|---|
| Cyclin D1 Antibody | CST | 55506 |
| Goat anti-Rabbit IRDye800 | Licor | 926-32211 |
| Goat anti-Mouse IRDye680 | Licor | 926-68070 |
| Goat anti-Human IgG (H+L) Cross-Adsorbed Secondary Antibody, Alexa Fluor™ 568 | Thermo Fisher | A-21090 |
| Goat anti-Rabbit IgG (H+L) Cross-Adsorbed Secondary Antibody, Alexa Fluor™ 488 | Thermo Fisher | 35553 |
| **In Vitro Binding Assays** | | |
| Pierce™ Streptavidin Coated Plates, 96-well black | Thermo Fisher | 15119 |
| Tween-20 | Sigma | P1379 |
| Recombinant Biotinylated HER2 AviTag | ACRO Biosystems | HE2H82E225UG |
| Recombinant Biotinylated CD36 AviTag | ACRO Biosystems | CD6H82E925UG |
| Recombinant Biotinylated INSR AviTag | ACRO Biosystems | INR-H82E6-25ug |
| Pierce™ Bovine Serum Albumin, Biotinylated | Thermo Fisher | 29130 |
| **siRNA Knockdown (KD) Reagents** | | |
| Opti-MEM™ I Reduced Serum Medium | Gibco | 31985062 |
| Lipofectamine™ 3000 Transfection Reagent | Invitrogen | L3000001 |
| Lipofectamine™ RNAiMAX Transfection Reagent | Invitrogen | 13778150 |
| ON-TARGETplus Non-targeting Pool scrambled | Dharmacon | D-001810-10 |
| GAL1 Human ON-TARGETplus siRNA SMARTPool | Dharmacon | L-011718-00-0005 |
| GAL3 Human ON-TARGETplus siRNA SMARTPool | Dharmacon | L-010606-00-0005 |
| PTPRF Human ON-TARGETplus siRNA SMARTPool | Dharmacon | L-008375-00-0020 |
| **Cell Proliferation Assays** | | |
| 96 Well White Polystyrene Microplate, TC treated | Corning | CLS3903 |
| RealTime-Glo™ MT Cell Viability Assay | Promega | G9711 |
| Trypan Blue Solution, 0.4% | Gibco | 15250061 |
| GB1107 | MedChemExpress | HY-114409 |
| **GB1211 Synthesis** | | |
| GB1211 | This work | |
| Copper(I) iodide | Sigma Aldrich | 03140 |
| tris((1-benzyl-4-triazolyl)methyl)amine | Ambeed | *A237566* |
| Cesium fluoride | Sigma Aldrich | 198323 |
| Triethylamine | Sigma Aldrich | 471283 |
| Sodium methoxide | Sigma Aldrich | 164992 |
| Dowex 50W X8 | Sigma Aldrich | 44519 |
| **Immunofluorescence** | | |
| Paraformaldehyde 16% | Thermo Fisher | 28906 |

| | | |
|---|---|---|
| Triton X-100 10% | Thermo Fisher | 85111 |
| Nunc™ Lab-Tek™ II Chambered Coverglass, 8-well | Thermo Fisher | 155409 |
| Poly-L-Lysine, 0.01% | Sigma | P4707 |
| DAPI | Thermo Fisher | 62248 |
| **Proximity Ligation Assay** | | |
| 96 Well glass bottom plate | Cellvis | P96-1.5H-N |
| Duolink® In Situ PLA® Probe Anti-Human PLUS | Sigma | DUO92020 |
| Duolink® In Situ PLA® Probe Anti-Rabbit MINUS | Sigma | DUO92005 |
| Duolink® In Situ Detection Reagents Red | Sigma | DUO92008 |

#### Cell Culture

All cells were maintained in a humidified incubator at 37 °C with 5% $CO_2$. All media was supplemented with 1% Penicillin-Streptomycin (final concentration: 100 U/mL penicillin, 100 µg/mL streptomycin). AU565, BT474, BT474 clone 5, HCC1419, HCC1954, and ZR-75-1 cells were cultured in RPMI-1640 medium supplemented with 10% FBS. MDA-MB-231, MDA-MB-361, and MDA-MB-453 cells were cultured in Leibovitz's L-15 medium supplemented with 10% FBS. MCF-7 and JIMT1 cells were cultured in DMEM supplemented with 10% FBS. MCF 10A cells cultured MEGM (Lonza) supplemented with 0.4% bovine pituitary extract (BPE), 10 ng/mL human epidermal growth factor (hEGF), 0.5 µg/mL hydrocortisone, 100 ng/mL Cholera Toxin, and 10 µg/mL insulin. Cells were routinely passaged upon reaching 70-90% confluency using TrypLE solution.

#### Preparation of Antibody-Catalyst Conjugates

500 µg of Human IgG isotype control or human anti-HER2 Trastuzumab biosimilar was first buffer exchanged using 10 kDa Amicon Ultra-Centrifugal Filters into 0.2 M Carbonate/BiCarbonate buffer, pH 9.2. Separately, a 5 mM solution of Ir catalyst DBCO was prepared in DMSO. 20 µl of this solution was then reacted with 10 µl of a 10 mM solution of azido-$PEG_{12}$-NHS ester in DMSO according to the scheme below. After incubating in the dark for 1 hour, 15 µl of this mixture was then reacted with 500 µg of buffer-exchanged antibody in a total volume of 250 µL and allowed to incubate with end-over-end rotation for 1 hour protected from light.

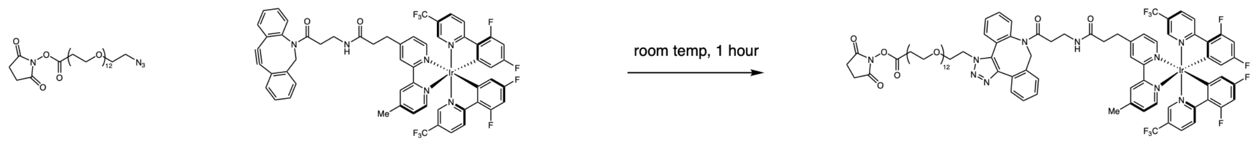

*azido-PEG12-NHS* *DBCO-Ir* *Ir-PEG12-NHS*

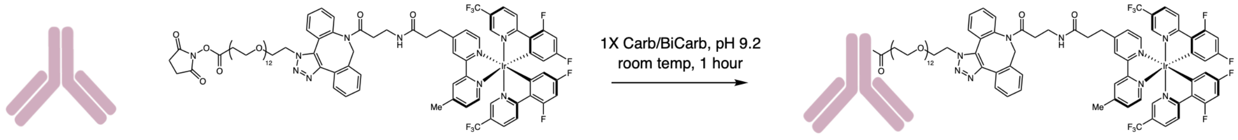

*antibody (500 µg)* *Ir-PEG12-NHS (10 eq.)* *Ir-conjugated antibody*

Crude conjugation reactions were then purified using 2 mL Zeba™ Spin Desalting Columns, 40K MWCO after equilibrating in 1X DPBS. Purified conjugated antibodies were first characterized by Pierce™ 660nm Protein Assay Reagent versus a bovine serum albumin standard curve. Catalyst loading was determined by comparing the absorbance of respective conjugates at 390 nm vs a Ir catalyst DBCO standard curve in PBS. Conjugation reactions typically yielded catalyst loading rates of ~1-3 catalysts per antibody.

#### Proximity labeling on AU565 Cells: Western Blot / Surface Staining

*Unless specified, all pelleting and washing of cells were performed at 300 x g for 4 min at 4 °C.* AU565 cells were dissociated from 10 cm culture dishes via 15 min incubation with TrypLE Express at 37 °C and quenched with complete RPMI. Cells were pooled together and resuspended in cold DPBS at 5 million cells/mL and then transferred in 1 mL aliquots to 1.7 mL Eppendorf Axygen Maxymum Recovery tubes. 10 µg of Isotype-Ir or Trastuzumab-Ir were then added to respective tubes and incubated on a rotator for 1 hour at 4 °C. After incubation, the cells were pelleted to remove the supernatant, washed twice with 1 mL cold DPBS, and resuspended in 1 mL of cold DPBS containing 500 µM biotin-PEG$_3$-diazirine (**S1**) (prepared from 500 mM stock in DMSO). The samples were then immediately placed in a 4 mL vial holder inside the m2 photoreactor and irradiated with blue light for 5 minutes in a 4 °C cold room. After irradiation, the cells were pelleted to remove the supernatant and washed twice with 1 mL cold DPBS. For each sample, a 100 µl aliquot (~500,000 cells) was transferred to a new tube and combined with 1 µl Streptavidin, Alexa Fluor™ 555 Conjugate and incubated for 30 minutes on ice protected from light. Cells were pelleted and washed 3x and resuspended in cold 100 µl DPBS. Samples were then transferred to black 96 well plates and fluorescence was measured using a BioTek plate reader (excitation 553 nm, emission 568 nm). The remainder of each photolabeled sample (900 µl, ~4,500,000 cells) was pelleted and then resuspended in 1 mL of membrane permeabilization buffer (MEM-PER Plus Membrane Fractionation Kit) containing 1X protease inhibitors and incubated for 20 min at room temperature with rotation. The samples were centrifuged at 16,000 x *g* for 15 min at 4 °C. The supernatant was removed, and the pellet was resuspended in 300 µL 1X RIPA containing 1% SDS and 1X protease inhibitors. The samples were placed on ice and sonicated using a Branson tip sonicator to break up the membrane pellet once for 5s at 35% intensity and heated for 5 min at 95 °C. The samples were diluted to 1.0 mL with 1X RIPA, transferred to 2.0 mL tubes, and sonicated an additional time for 5s at 35% intensity. The lysates were then centrifuged at 16,000 x *g* to pellet any insoluble material, and the supernatants transferred to a new tube and protein concentration quantified by BCA. 100 µg of each lysate was set aside as “input” for western blot analysis. For streptavidin bead enrichment, the remaining cell lysate (~1 mg) was added to a 1.7 mL Axygen tube containing 100 µL of Pierce streptavidin magnetic beads that were pre-washed twice with 1 mL RIPA. The samples were incubated overnight with rotation at 4 °C, and the beads were pelleted on a magnetic rack. The supernatant was removed, and the beads were washed 3x with 1 mL 1% SDS in DPBS, 3x with 1 mL 1M NaCl in DPBS, and 3x with 1 mL 10% EtOH in DPBS. The beads were then pelleted to remove supernatant and resuspended in 40 µL of elution buffer (6 M Urea, 2 M thiourea, 30 mM biotin, 2% SDS in DPBS, 25% 1x Laemmli with 2-mercaptoethanol, pH=11.5). The samples were heated at 95 °C for 15 minutes with shaking at 700 rpm. The beads were pelleted on a magnetic rack, and the supernatant collected. 20 µL of this material and 10 μg of each input (~1%) were loaded on a 4-20% Criterion TGX Precast SDS-PAGE Gel. Gels were run in freshly prepared tris-glycine running buffer for 1 h at 160 V using a Criterion Cell and Powerpac (BioRad,) For western analysis, blots were probed with rabbit Anti-HER2 primary (1:1000 dilution in 5% BSA in TBST, Cell Signaling Technologies) overnight at 4 °C. The membrane was washed 3x with TBST (5 min) and and incubated in blocking buffer with 1:5000 secondary antibody (Goat anti-rabbit IR Dye 800 CW) for 1 h at room temperature. The membrane was washed 3x with TBST (5 min) before imaging using an Odyssey CLx (LiCor, 9140).

#### Western Blot

Gel electrophoresis was performed using a Bio-Rad Criterion Vertical Electrophoresis Cell tank, BioRad PowerPac Basic Power Supply, and Criterion TGX tris-glycine polyacrylamide gel cassettes (SDS/Tris) using tris-glycine-SDS running buffer (25 mM Tris, 200 mM glycine, 35 mM SDS). Gels were transferred from precast cassettes to nitrocellulose membranes (0.45 μm, BioRad 1620115) using a BioRad Criterion blotter apparatus. The membranes were then blocked in 5% bovine serum album (BSA) in Tris-buffered saline + 0.05% Tween20 (TBST) and incubated for 1 hour at room temperature with agitation. The blocking solution was then decanted and replaced with a primary antibody solution in blocking buffer and incubated overnight at 4 °C with gentle rocking. Membranes were then washed with 1X TBST (3 x 5 min) before incubation with a secondary antibody solution followed by imaging via Licor Odyssey CLx scanner Densitometry was performed using Image Studio Lite V. 5.2 (Licor) or Fiji.

#### HER2 *μ*Map Proximity Labeling Workflow (Proteomics)

Cell lines were seeded into 10 cm dishes with three biological replicates for each condition (isotype-Ir and trastuzumab-Ir) and grown to ~90% confluency on the day of labeling. Cells were dissociated from culture dishes via 15 min incubation with TrypLE Express at 37 °C and quenched with complete media. Cells were pooled together and resuspended in cold DPBS then transferred in triplicate 1 mL aliquots for each condition in 1.7 mL

Eppendorf Axygen Maxymum Recovery tubes. 10 µg of Isotype-Ir or Trastuzumab-Ir were then added to respective tubes and incubated on a rotator for 1 hour at 4 °C. After incubation, the cells were pelleted to remove the supernatant, washed twice with 1 mL cold DPBS, and resuspended in 1 mL of cold DPBS containing 500 µM biotin-$PEG_3$-diazirine (**S1**) (prepared from 500 mM stock in DMSO). The samples were then immediately placed in a 4 mL vial holder inside the m2 photoreactor and irradiated with blue light for 5 minutes in a 4 ºC cold room. After irradiation, the cells were pelleted to remove the supernatant and washed twice with 1 mL cold DPBS. The pellet was resuspended in 1 mL of membrane permeabilization buffer (MEM-PER Plus Membrane Fractionation Kit) containing 1X protease inhibitors and incubated for 20 min at room temperature with rotation. The samples were centrifuged at 16,000 x *g* for 15 min at 4 °C. The supernatant was removed, and the pellet was resuspended in 300 µL 1X RIPA containing 1% SDS and 1X protease inhibitors. The samples were placed on ice and sonicated using a Branson tip sonicator to break up the membrane pellet once for 5s at 35% intensity and heated for 5 min at 95 °C. The samples were diluted to 1.0 mL with 1X RIPA, transferred to 2.0 mL tubes, and sonicated an additional time for 5s at 35% intensity. The lysates were then centrifuged at 16,000 x *g* to pellet any insoluble material, and the supernatants transferred to a new tube. For streptavidin bead enrichment, 1 mg of membrane cell lysate was added to a 1.7 mL Axygen tube containing 100 µL of Pierce streptavidin magnetic beads that were pre-washed twice with 1 mL RIPA buffer. The samples were incubated overnight on a rotisserie at 4 ºC, and the beads were pelleted on a magnetic rack. The supernatant was removed, and the beads were washed 3x with 1 mL 1% SDS in DPBS, 3x with 1 mL 1M NaCl in DPBS, 3x with 1 mL 10% EtOH in DPBS, and 3x with 1 mL 100 mM $NH_4HCO_3$. The beads were then resuspended in 500 µL 6 M urea in DPBS and 25 µL of 200 mM DTT in 25 mM $NH_4HCO_3$ was added. The beads were then incubated at 55 ºC for 30 minutes on a rotator. Subsequently, 30 µL 500 mM iodoacetamide (IAA) in 25 mM $NH_4HCO_3$ was added and incubated for 30 min on a rotator at room temperature in the dark. The supernatant was removed, and the beads were washed 3x with 0.5 mL DPBS followed by 3x with 50 mM $NH_4HCO_3$. Beads were then resuspended in 40 $\mu$L 50 mM Ammonium Bicarbonate buffer containing 1 $\mu$g MS-grade trypsin per sample and digested at 37 °C overnight with rotation. Trypsin was then quenched with 1 μL Optima LC/MS Formic Acid and supernatants were passed through a 0.22-micron spin filter column (CoStar, 98231-UT-1) before being transferred to an LC/MS vial for proteomics analysis.

#### Quantitative Mass Spectrometry Proteomics and Data Analysis

Label-free, data-independent acquisition (DIA) proteomics was performed on a Bruker TimsTOF Pro 2 connected to a nanoElute LC. For each sample, ~100 ng of protein was injected onto a trap column (C18 Pepmap, 5 μM particle size, 5 mm length, 300 μM internal diameter), followed by separation via an analytical column (C18 ReproSil AQ, 1.9 μM particle size, 100 mm length, 75 μM internal diameter). Peptides were eluted via an acetonitrile/water gradient at a column temperature of 40 °C (buffer A = 0.1% formic acid/water, buffer B = 0.1% formic acid/acetonitrile; flow rate 0.5 μL/min; gradient: start at 2% B, then increase to 35% B over 20 min, increase to 95% B over 0.5 min, hold at 95% for 2.25 min). Scans were performed in positive ion, dia-PASEF mode over a m/z range of 100-1700 with a ramp time of 100 ms, Accu. time of 100 ms, and a duty cycle of 100%, ramp rate of 9.43 Hz, MS averaging set to 1. Absolute thresholds were set to 5000 for mobility peaks and 10 for MS peaks. Data were collected with Bruker Compass HyStar v6.2. Raw data (.d files) were then processed via DIA-NN 1.8.1 via the following parameters: trypsin/P digestion, 3 missed cleavages, 3 max. variable modifications, N-term M excision, Ox(M), Ac(N-term) and C-carbamidomethylation, peptide length range of 7-30, precursor charge range 1-4, m/z range 300-1800, fragment ion range 200-1800, Mass accuracy and MS accuracy both set to 10, precursor FDR set to 1%. Within the DIA-NN algorithm, the following settings are applied: “Use isotopologues”, “MBR” (match between runs), “No shared spectra”, “Heuristic protein inference”. A spectral library was utilized which was generated in DIA-NN from all known human proteins (In-Silico spectral library – generated in DIA-NN via FASTA of Uniprot human proteome UP000005640 – options selected were “FASTA digest for library-free search/library generation” and “Deep learning-based spectra, RTs and IMs prediction”, other parameters same as described above). After processing, the resulting matrix.pg files were analyzed in Perseus (v 2.0.7.0)[71], where intensities are inputted as “main” while the other descriptors are listed as “categorical”. Intensities were transformed by log base 2, and data was annotated to group by the appropriate condition. Missing values were then replaced from a normal distribution (width = 0.3, downshift = 1.8, separately for each column). Normalization was performed via median subtraction, and a volcano plot was generated utilizing a t-test for statistical significance. Resulting volcano plots were plotted in GraphPad Prism 10 for final figures. Gene ontology analysis was performed on Metascape v3.5.

#### In Vitro Trastuzumab Recombinant Binding Assay

96-well black streptavidin coated plates were warmed to room temperature and washed once with 150 $\mu$l 1X Tris-buffered saline + 0.05% Tween20 (TBST). Recombinant proteins were diluted to 1 $\mu$g/mL solution in blocking buffer (3% BSA in TBST). 100 $\mu$L of this solution was added to each well (~100 ng/well) and incubated for 1 hour at room temperature with gentle shaking. The solution was then removed and each well was washed 3x with 150 $\mu$l TBST. To respective wells, a range of concentrations of either Trastuzumab or Human isotype control (1 $\mu$g - ~15 ng in 100 $\mu$l volume blocking buffer) was added and the plate was incubated for 1 hour at room temperature with gentle shaking. The solution was then removed and each well was washed 3x with 150 $\mu$l TBST, and 100 $\mu$l of a secondary antibody solution (Goat anti-Human AlexaFluor 568, 1:1000 in blocking buffer) was added to each well followed by incubation for 1 hour at room temperature protected from light with gentle shaking. The solution was then removed and each well was washed 3x with 150 $\mu$l TBST and the fluorescence of each well was measured with BioTek plate reader at 578 nm excitation and 603 nm emission.

#### siRNA Knockdown and Cell Viability Assays (HCC1954)

HCC1954 cells were seeded into white-walled, clear bottom 96-well plates at 1,000 cells per well and allowed to recover overnight. To appropriate wells, 0.1 mL of a 10 $\mu$g/mL solution of trastuzumab was prepared in complete RPMI and added. siRNA (non-targeted scrambled or target) transfection mixes were prepared by combining 10 $\mu$L of each siRNA (10 $\mu$M stock in water, 100 pmol total) with 117 $\mu$L OptiMEM media in one tube while another tube was separately filled with 117 $\mu$L OptiMEM media and 7.5 $\mu$L Lipofectamine 3000 transfection reagent. The two transfection mixes were then combined and incubated at room temperature for 5 minutes. 10 $\mu$l of each appropriate mix was then added dropwise to respective treatment wells and incubated at 37 °C, 5% $CO_2$ for 48 h. Another redose of trastuzumab and siRNA was given to respective wells and incubated for another 48 h (day 4 time point). At day 2 and day 4 timepoints, media was removed from each well and replaced with 100 $\mu$L of a 1X solution of RealTime-Glo MT Cell Viability Assay reagent in complete RPMI containing MT Cell Viability Substrate and NanoLuc Enzyme. Cells were incubated at 37 °C, 5% $CO_2$ and luminescence was then measured using a BioTek Gen5 v2.05 plate reader. Viability was calculated by comparing to untreated cells receiving vehicle (media).

#### GB1211 Synthesis

(2R,3R,4S,5R,6R)-2-((5-bromopyridin-3-yl)thio)-6-(hydroxymethyl)-4-(4-(3,4,5-trifluoro-phenyl)-1H-1,2,3-triazol-1-yl)tetrahydro-2H-pyran-3,5-diol (GB1211/Selvigaltin)

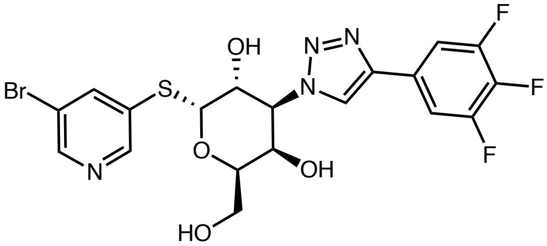

The synthetic preparation of GB1211 is adapted from the published route (*Journal of Medicinal Chemistry*, **2022**, 26;65(19):12626-38). To a solution of 5-bromopyridin-3-yl 2,4,6-tri-*O*-acetyl-3-azido-3-deoxy-l-thio-a-d-galactopyranoside (1.05 g, 2.09 mmol) in THF (25 mL) was added trimethyl-[2-(3,4,5-trifluorophenyl)ethynyl]silane (714 mg, 314 mmol), CuI (40 mg, 0.21 mmol), tris((1-benzyl-4-triazolyl)methyl)amine (TBTA) (111 mg, 0.12 mmol), CsF (475 mg, 3.13 mmol) and triethylamine (1.8 mL). The reaction vessel was purged 3 times with nitrogen. The reaction mixture was stirred at 50 °C for 2

h. The mixture was filtered over Celite and washed with EtOAc (2 x 50 mL), The filtrate was concentrated in vacuo to afford the crude cycloaddition product, which was used for next step directly without further purification. The crude product was dissolved in MeOH (20 mL) followed by addition of NaOMe (16.0 mg, 0.3 mmol). The mixture was stirred at room temperature for 1 h, after which no more starting material was detected by UPLC analysis, followed by acidification to pH 5–6 using Dowex 50W X8 hydrogen form. The reaction mixture was filtered, washed with MeOH (40 mL) and concentrated in vacuo to afford the crude product, which was purified by preparative HPLC (RediSep C18 column, 15% to 80% MeCN in $H_2O$ with 0.1% formic acid) to provide selvigaltin as a beige solid (700 mg, 1.31 mmol, 63% yield).

**$^1$H NMR (500 MHz, $CD_3OD$)** δ 8.70 (d, *J* = 1.9 Hz, 1H), 8.59 (d, J = 2.1 Hz, 1H), 8.57 (s, 1H), 8.34 (t, *J* = 1.9 Hz, 1H), 7.64 (dd, J = 9.0, 6.5 Hz, 2H), 5.93 (d, *J* = 5.2 Hz, 1H), 5.00 (dd, J = 11.2, 2.9 Hz, 1H), 4.98 (dd, *J* = 11.4, 5.2 Hz, 1H), 4.50 (dt, *J* = 6.0, 0.9 Hz, 1H), 4.20 (dd, *J* = 2.6, 1.0 Hz, 1H), 3.70 (dd, *J* = 11.5, 5.4 Hz, 1H), 3.70 (dd, J = 11.5, 6.3 Hz, 1H).

**$^{13}$C NMR (126 MHz, $CD_3OD$)** δ 151.0, 145.0, 144.0, 135.2, 122.8, 121.8, 110.8, 91.0, 73.4, 69.9, 66.9, 65.4, 62.1.

The spectroscopic data match the reported literature values.

#### Galectin Inhibition and Trastuzumab Cell Viability Assays

Each tested cell line was seeded into white-walled, clear bottom 96-well plates at 1,000 cells per well and allowed to recover overnight. Initial testing of GB1107 and GB1211 was performed by first preparing 50 mg/mL stock solutions of each compound in DMSO. To appropriate wells, 0.1 mL of complete RPMI containing either 0.1% DMSO or various dilutions of each compound (50 $\mu$g/mL, 25 $\mu$g/mL, etc) was added and the plates were incubated at 37 °C, 5% $CO_2$ for 3 days followed by viability measurement using the RealTime-Glo MT Cell Viability Assay. For combination treatments in cell lines, 0.1 mL of complete RPMI containing either 0.1% DMSO, 10 $\mu$g/mL of trastuzumab, 10 $\mu$g/mL GB1211, or combination was added to appropriate wells and the plates incubated at 37

°C, 5% $CO_2$ for 48 h. Another redose was given to respective wells every 48 h. At each desired timepoint, viability was then measured using the RealTime-Glo MT Cell Viability Assay kit compared to vehicle control wells.

#### Immunofluorescence Microscopy

AU565 and HCC1954 cells were seeded into glass-bottom 8-well chamber slides pre-coated with 0.01% poly-L-lysine and allowed to recover at 37 °C, 5% $CO_2$ for 48 h. Cells were then fixed with 4% paraformaldehyde in DPBS for 15 minutes at room temperature, washed once with DPBS, and then permeabilized with 0.1% Triton X-100 in DPBS for 10 minutes. Samples were then incubated at room temperature for 1 hour in blocking buffer (3% BSA, 0.05% Tween20 in DPBS). A primary antibody solution (Trastuzumab Biorad, 4D5-8, 10 $\mu$g/mL; GAL1 CST D608T, 1:50; GAL3 CST D4I2R 1:500) was then prepared in blocking buffer and applied for at least 1 hour at room temperature. Samples were then washed 3X with 0.05% Tw20 in DPBS and stained with a secondary antibody solution in Blocking buffer for 1 hour at room temperature. DAPI (1:10000) was used for nuclear staining, goat anti-rabbit Alexa Fluor 488 (1:1000) and goat anti-human Alexa Fluor 568 (1:1000) were used to target specific primary antibodies. Finally, samples were washed 3X with 0.05% Tw20 in DPBS and stored in DPBS before imaging. Samples were imaged using NIS Elements AR v4.60.00 software on a Nikon A1R-Si HD Confocal Microscope. Image processing was performed with ImageJ2/FIJI Mac OS X software.

#### Proximity Ligation Assay

Cells were seeded into glass-bottom 96-well plates pre-coated with 0.01% poly-L-lysine and allowed to recover at 37 °C, 5% $CO_2$ for 24-48 h. Cells were then fixed with 4% paraformaldehyde in DPBS for 15 minutes at room temperature, washed once with DPBS, and then permeabilized with 0.1% Triton X-100 in DPBS for 10 minutes. Samples were then incubated at room temperature for 1 hour in blocking buffer (3% BSA, 0.05% Tween20 in DPBS). A primary antibody solution (Trastuzumab Biorad, 4D5-8, 10 $\mu$g/mL; GAL1 CST D608T, 1:50; GAL3 CST D4I2R 1:500) was then prepared and added to each well. Plates were incubated for at least 1 hour at room temperature. Each well was then washed 3x with DPBS and then incubated with an appropriate PLA probe solution (Goat anti-Human PLUS or Goat anti-Rabbit MINUS) that was diluted 1:5 in 3% BSA in DPBS. Plates were incubated at 37 °C for at least an hour, followed by a 5 minute incubation with 1X Wash Buffer A. Wells were then incubated with ligase (1:40 in 1X ligation buffer) and incubated at 37 °C for 30 minutes. After 2x5 minute washes with 1X Wash Buffer A, wells were incubated with polymerase (1:80 in 1X amplification buffer) at 37 °C for 100 minutes. Wells were washed twice with 1X Wash Buffer B for 10 minutes followed by one wash with 0.01X Wash Buffer B. After airdrying wells for 5 minutes, they were then covered with a minimal amount of Duolink In Situ Mounting Medium. Samples were imaged using NIS Elements AR v4.60.00 software on a Nikon A1R-Si HD Confocal Microscope. Image processing was performed with ImageJ2/FIJI Mac OS X software.

#### Trastuzumab Cell Binding Assay

HCC1954 cells were grown in either 10 $\mu$g/mL GB1211 or 0.1% DMSO in complete RPMI for 48 h, followed by another redose and incubation for 48 h. Cells were then harvested and counted and resuspended in cold DPBS. $1x10^6$ cells was then transferred in triplicate 1 mL aliquots for each condition in 1.7 mL Eppendorf Axygen Maxymum Recovery tubes. 10 µg of human isotype control or trastuzumab were then added to respective tubes and incubated on a rotator for 1 hour at 4 °C. After incubation, the cells were pelleted to remove the supernatant, washed twice with 1 mL cold DPBS, and resuspended in 1 mL cold DPBS containing Goat anti-Human AlexaFluor 568 (1:500). After incubation at 4 °C for 1 hour with rotation, cells were again pelleted and washed twice with 1 mL cold DPBS. Cells were pelleted and resuspended in 500 µl DPBS and 100 µl from each binding reaction was transferred to a 96-well black plate and fluorescence of each well was measured with a BioTek plate reader at 578 nm excitation and 603 nm emission.

#### Global Proteomics Sample Preparation

HCC1954 cells were cultured in triplicate 10 cm dishes for each condition (DMSO, Tz, GB1211, or Tz + GB1211). 10 mL of complete RPMI containing either 0.1% DMSO, 10 µg/mL of trastuzumab, 10 µg/mL GB1211, or combination was added to appropriate dishes and incubated at 37 °C, 5% $CO_2$ for 48 h. Another redose was given and the dishes incubated another 48 h. Dishes were washed once in 1X DPBS and cells were harvested by gentle scraping. Cell pellets were then resuspended in 1X RIPA and sonicated for 5s at 35% intensity. The lysates were then centrifuged at 16,000 x *g* to pellet any insoluble material, and the supernatants transferred to a new tube. Protein concentrations of lysates were determined via BCA. For each replicate, Add 250 µg of whole cell lysate was diluted with RIPA to 250 µL. DTT was then added to a final concentration of 10 mM and lysate was heated at 95 °C for 5 minutes. Iodoacetamide was then added to a final concentration of 15 mM and tubes were incubated in the dark with rotation for 30 minutes. DTT was then added to obtain a final concentration of 20 mM to quench IAA and incubated at room temperature for 15 minutes with rotation. 10 µg of reduced and alkylated lysates was diluted to a final volume of 48 µL in reconstitution solution (50 mM HEPES pH 8, 1% SDS, 1% Tx100, 1% NP-40, 1%Tw20, 1% deoxycholate, 5 mM EDTA, 50 mM NaCl, 5 mM DTT, 1% glycerol in LC/MS grade water). 2 µL of a prepared 50 µg/µL seramage bead stock, was then added to each tube, followed by 50 *µ*L ethanol. Tubes were then incubated at 24 °C for 5 min at with shaking at 1,000 r.p.m. Beads were then pelleted on a magnetic rack and unbound supernatant was discarded. Beads were then washed three times with 200 *µ*L 80% ethanol and supernatant was completely removed. Beads were then resuspended in 50 mM Ammonium Bicarbonate and 1 *µ*g trypsin was added to each tube and sonicated for 30 s in a water bath to fully disaggregate the beads, followed by incubation for 18 h at 37 °C with rotation. Trypsin was then quenched with 1 *µ*L Optima LC/MS Formic Acid and supernatants were passed through a 0.22-micron spin filter column (CoStar, 98231-UT-1) before being transferred to an LC/MS vial for proteomics analysis.

#### PTPRF Knockdown and Cell Viability Assays

Target cell lines were seeded into six 10 cm dishes and allowed to recover overnight. siRNA (non-targeted scrambled or PTPRF) transfection mixes were prepared by combining 60 μL of siRNA (10 μM stock, 600 pmol total) with 2.3 mL OptiMEM media in one tube while another tube was separately filled with 2.3 mL OptiMEM media and 150 μL RNAiMAX lipofection transfection reagent. The two transfection mixes were then combined and incubated at room temperature for 5 minutes. 1.6 mL of each appropriate mix was then added dropwise to triplicate dishes and incubated at 37 °C, 5% $CO_2$ for 48 h. Another redose of siRNA was performed and cells were incubated for another 48 h. This was repeated for 6 days total. Media and cells from each dish were then collected by gentle scraping and viable cells per mL were determined by mixing 10 μl cell suspension with 10 μL 0.4% Trypan Blue and analyzed using a Countess 3 Automated Cell Counter (Invitrogen). The remaining cell suspension was then pelleted and lysates were prepared for western blot analysis.

#### Structures of additional reagents used for experiments

S1 was prepared as described previously (*Science*, **2020**, *367*(6482), 1091-1097). Ir DBCO catalyst (S2) was prepared as described previously (*J. Am. Chem Soc.*, **2022**, *144* (51) 23633-23641).

Biotin-$PEG_3$-diazirine (S1)

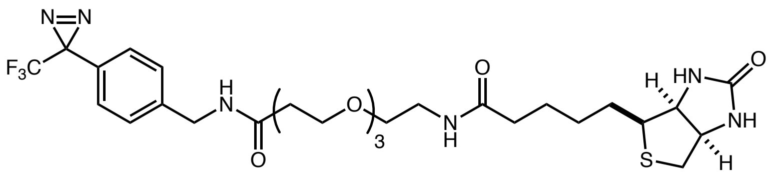

Ir DBCO Catalyst (S2)

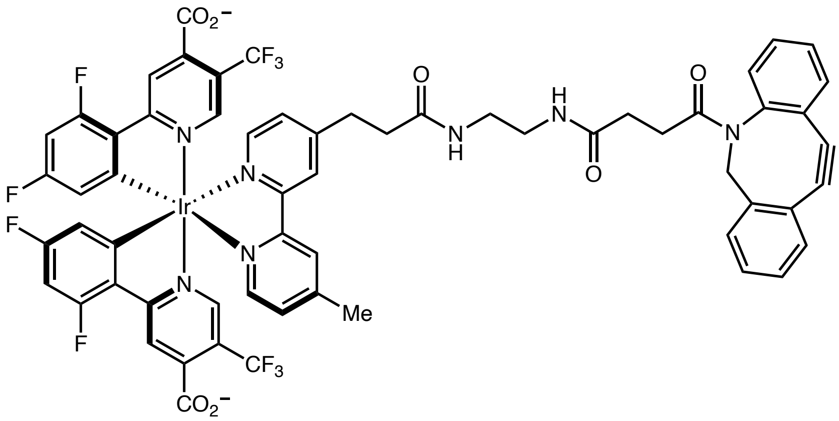
CO2−
CF3
F
F
N
N
Ir
N
N
F
F
O
N
H
H
N
O
O
N
Me
CF3
CO2−
